# The neuron mixer and its impact on human brain dynamics

**DOI:** 10.1101/2023.01.05.522833

**Authors:** Charlotte E. Luff, Robert Peach, Emma-Jane Mallas, Edward Rhodes, Felix Laumann, Edward S. Boyden, David J. Sharp, Mauricio Barahona, Nir Grossman

**Affiliations:** Department of Brain Sciences, Imperial College London; London, United Kingdom; UK Dementia Research Institute, Imperial College London; London, United Kingdom; Department of Neurology, University Hospital Würzburg; Würzburg, Germany; UK Dementia Research Institute, Care Research & Technology Centre; London, United Kingdom; Department of Mathematics, Imperial College London; London, United Kingdom; Department of Brain and Cognitive Sciences, Massachusetts Institute of Technology; Cambridge, USA; Howard Hughes Medical Institute; Chevy Chase, USA; Centre for Injury Studies, Imperial College London; London, United Kingdom

## Abstract

A signal mixer made of a transistor or a diode facilitates rich computation, which has been the building block of modern telecommunications. The mixing produces new signals at the sum and difference frequencies of the input, thereby enabling powerful operations such as frequency conversion (aka heterodyning), phase detection, and multiplexing. Here, we report that a neural cell is also a signal mixer. We found through *ex-vivo* and *in-vivo* whole-cell measurements that neurons mix exogenous (controlled) and endogenous (spontaneous) subthreshold membrane potential oscillations, producing new oscillation frequencies. We show, using pharmacological manipulation, that the neural mixing originates in the voltage-gated ion channels. Furthermore, we demonstrate that the neural mixing dynamic is evident in human brain activity and is associated with cognitive functions. We found that the human electroencephalogram (EEG) displays distinct clusters of local and inter-region mixing interactions. By quantifying the strength of these interactions before a task, we show that converting the salient posterior alpha-beta oscillations into gamma-band oscillations regulates the visual attention state. Neural circuit oscillations have been observed in nearly every cognitive domain and species, and abnormal spectra of neural oscillations have been found in almost all brain disorders. Signal mixing enables individual neurons to actively sculpt the spectrum of their circuit oscillations and utilize them for computational operations, which have only been seen in modern telecommunication until now.

## Introduction

A mixer is a nonlinear device capable of multiplying signals to produce new frequencies, such as the difference and sum of the original input frequencies. A signal mixer made of a transistor, or a diode, has been the building block of modern telecommunication, facilitating the critical conversion to/from higher frequency bands where transmission efficiency is high (aka heterodyning), decoding phase information, and combining multiple signals into one data stream (aka multiplexing) (Goldsmith, 2005; Protopopov, 2010).

Various neuroscientific studies have reported evidence of a multiplication operation in individual neurons, attributed to their need to implement coincidence detection, i.e., the joint probability of two statistically independent events is the product of probabilities of the individual events (Srinivasan & Bernard, 1976). Neural coincidence detection has been observed in diverse tasks, such as the transformation of eye-centric into head-centric coordinates (Andersen et al., 1985), localization of sound (Peña & Konishi, 2001), combination of multisensory signals (Haag et al., 2010; Huston & Krapp, 2009) and detection of visual motion (Hatsopoulos et al., 1995; Von Hassenstein & Reichardt, 1956). The biophysical underpinning of the multiplication operation has been linked to the nonlinear transfer function of synaptic currents, including log-exp transformation of coincident synaptic excitatory and inhibitory events (Gabbiani et al., 2002), concurrent synaptic excitation and release from shunting inhibition events (Groschner et al., 2022), as well as to the voltage-dependent inactivation of the NMDA receptors (Lavzin et al., 2012; Poleg-Polsky & Diamond, 2016).

Recent studies have demonstrated the frequency mixing of neural oscillations in rodents’ brains (also known as signal mixing or spectral mixing but distinct from cross-frequency coupling). Ahrens et al. used local field potential (LFP) recordings in rodents, to show that mechanically evoked neural oscillations in the vibrissae can mix internally, and with spontaneous oscillations induced by anaesthesia (Ahrens et al., 2002). Haufler et al. deployed a statistical phase analysis to demonstrate phase dependency between frequency mixing quadruplets (i.e., roots: *f*1, *f*2; products: Δ*f*, ∑*f*) in the spontaneous LFP activity of rodents in a state and region-dependent manner (Haufler & Paré, 2019). Using a computational model, it was proposed that the signal mixing could emerge from individual neurons’ nonlinear threshold depolarization characteristics (Ahrens et al., 2002; Kleinfeld and Mehta, 2006) (a linear response cannot lead to mixing).

To date, there has not been experimental evidence that an individual neuron can mix transmembrane potentials, and the contribution of this phenomenon to human brain activity has not been explored. Here, we use whole-cell patch clamp recording in mice, to experimentally demonstrate that individual neurons mix exogenous and endogenous transmembrane potential signals, and that the signal mixing originates in the nonlinear voltage-gated ion channel currents. Then, we apply novel phase computation statistics to electroencephalogram (EEG) recordings in human subjects, to demonstrate cognitively relevant frequency mixing in the human brain.

## Results

### Mixing of exogenous membrane potentials in individual neurons

We first examined whether neurons could mix exogenous (controlled) subthreshold membrane oscillations. We applied extracellular sinusoidal electric currents containing two different frequencies (*f*_1_ and *f*_2_), with a difference frequency (Δ*f*) within the normal range of neural activity, and recorded the induced transmembrane potentials in individual neurons *ex vivo* using whole-cell patch clamp recording (**Fig. 1A**). We focused on the subthreshold response of the neurons, because suprathreshold spikes at the difference frequency could also be induced by a summation of the applied fields, rather than mixing, i.e., neurons are firing due to polarisation at the applied frequencies periodically reaching the action potential threshold. We found that electrical stimulation with two sinusoids at frequencies within the normal range of neural activity (i.e., *f*_1_ = 47 Hz and *f*_2_ = 57 Hz), induced subthreshold membrane potential oscillations at their difference frequency (Δ*f* = 10 Hz) (**Fig. 1B**, note that the stimulation voltages mask the induced subthreshold oscillations at the input frequencies *f*_1_ and *f*_2_).

**Fig. 1.**
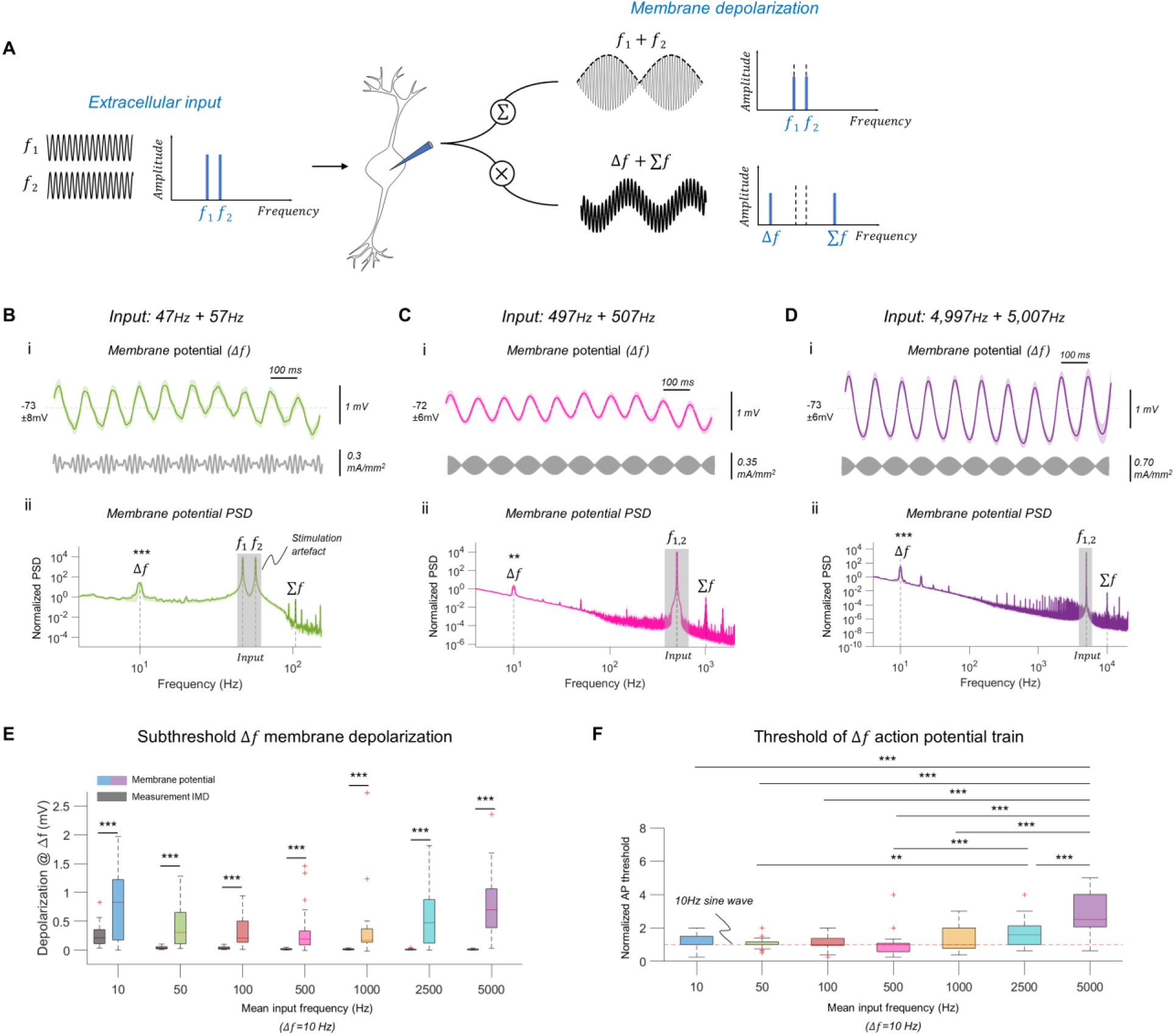
Mixing of exogenous membrane potentials in individual neurons *ex vivo*. (**A**) Neural mixing concept showing the subthreshold membrane transfer function of multi-frequency input with a conventional linear superposition (‘∑’), and the proposed nonlinear mixing via multiplication (‘X’). (**B**) (i) Top: Neural membrane potential during sinusoidal electrical stimulation with frequencies *f*_1_=47Hz + *f*_2_=57Hz (shown are mean ±SEM). Resting membrane potential mean ±SD is displayed. Raw membrane traces were filtered to remove stimulation artifact; n=21 cells. Bottom: Applied stimulation current waveform. (ii) Corresponding membrane potential’s power spectral density (PSD), mean ±SEM. Raw membrane traces were first filtered to remove offset. PSD values were normalized to endogenous PSD activity at 4Hz; n=21 cells. PSD at *f*_1_ and *f*_2_ is dominated by stimulation artifact. ***, p<0.0005, significant PSD peak, one tailed Wilcoxon signed rank test for zero median. (**C**) As in (B) but during stimulation with *f*_1_=497Hz + *f*_2_=597Hz; n=27 cells. **, p<0.005, one tailed Wilcoxon signed rank test for zero median. (**D**) As in (B) but during stimulation with *f*_1_=4997Hz + *f*_2_=5007Hz; n=29 cells. (**E**) Box plot showing root mean square (RMS) amplitude of the induced neural oscillation at Δ*f* (‘Membrane potential’) vs. the measurements’ IMD at Δ*f* (‘Measurement IMD’) across the range of stimulation frequencies. Traces were first filtered at Δ*f*. RMS values were baseline subtracted. n (IMD/Membrane potential) = 20/27 (10Hz); 20/21 (50Hz); 25/27 (100Hz); 26/27 (500Hz); 21/25 (1,000Hz); 29/29 (2,500Hz); 28/29 (5,000Hz) recordings/cells. ***, p < 0.0005, Wilcoxon rank sum test/two sample t-test. Current densities: 0.41±0.31 (10Hz); 0.30±0.24 (50Hz); 0.34±0.27 (100Hz); 0.34±0.22 (500Hz); 0.33±0.26 (1,000Hz); 0.45±0.34 (2,500Hz); 0.69±0.37 (5,000Hz) mA/mm^2^. (**F**) Normalized current threshold for action potential (AP) train at Δ*f* across the range of stimulation frequencies. Thresholds were normalized to threshold of a stimulation with 10Hz sine wave (horizontal red dashed line). *, comparisons survived Bonferroni correction (p-value=0.0071); **, p<0.005; ***, p<0.0005; repeated-measures ANOVA, post-hoc paired t-test. Boxplots: central mark, median; box edges, 25th and 75th percentiles; whiskers, extend up to 1.5x interquartile range box edges; ‘+’, datapoints outside this range. n = 22 (10Hz); 22 (50Hz); 21 (100Hz); 21 (500Hz); 20 (1,000Hz); 20 (2,500Hz); 19 (5,000Hz) cells.

The induction of a subthreshold membrane oscillation at the difference frequency was consistent across a wide range of stimulation frequencies, spanning three orders of magnitude. **Figure 1C** and **Figure 1D** show the subthreshold membrane oscillation induced by electrical stimulation with two sinusoids at frequencies in the upper boundary of neural activity (i.e., *f*_1_ = 497 Hz and *f*_2_ = 507 Hz, Δ*f*= 10 Hz) and far beyond the range of neural activity (i.e., *f*_1_ = 4.997 kHz and *f*_2_ = 5.007 kHz, Δ*f*= 10 Hz) as in temporal interference (TI) stimulation (Grossman et al., 2017; Violante et al., 2023), respectively. **Figure 1E** summarizes the induced oscillation amplitude at Δ*f* across this range of applied frequencies (tested against the measurement system’s intermodulation distortion, IMD at Δ*f* (Kasten et al., 2018), i.e., mixing products due to hardware nonlinearity measured in the same way but without brain slices). See **fig. S1** for representative unfiltered membrane potential recordings, and **fig. S2** for additional membrane potential traces. The amplitude of the induced Δ*f* oscillation was smaller when the stimulation was applied at kHz frequencies. Increasing the amplitude of the applied currents evoked action potential trains at Δ*f*, with a higher current density threshold at kHz frequencies (repeated-measures ANOVA F(5,135) = 22.3, p = 5e-17, **Fig. 1F**).

The membrane potential power spectra also showed peaks at the sum (∑*f*) and second harmonics (2*f*_1_, 2*f*_2_) of the applied frequencies (**Fig. 1B**-**D**, panel ii), as predicted by signal mixing. However, the low membrane oscillation amplitudes at those high frequencies were within the range of the measurement’s IMD, rendering these measurements inconclusive, except in the lowest stimulation frequency condition (i.e., *f*_1_ = 7 Hz and *f*_2_ = 17 Hz)-when the induced frequencies were within the range of normal neural activity (i.e., ∑*f* = 24 Hz, 2*f*_1_ = 14 Hz, 2*f*_2_ = 34 Hz). See **fig. S2** for all Δ*f* and ∑*f* membrane potential traces, **fig. S3** for statistical comparisons of the sum and harmonic frequencies, **table S1** for a summary of all investigated frequencies, and **tables S2 and S3** for all values and statistics for fig. 1. These results suggest that in addition to the difference frequency, neurons are capable of producing the sum frequency and harmonics of their membrane oscillations. Interestingly, we also found some significant subthreshold membrane oscillations at higher frequency mixing orders, including the second harmonic of the difference frequency (i.e., 2Δ*f* = 20 Hz; **fig. S3D**), but not at even higher order mixings (i.e., 2*f*_2_ – *f*_1_ and ∑*f* – 2Δ*f*; **fig. S3E-F)**, suggesting the neurons are capable of further mixing the mixing products, but that the mixing products at higher orders are potentially harder to distinguish from the measurement IMD (see **table S4** for all values and statistics for **fig. S3**).

### Origin of neuronal mixing characterized via pharmacological manipulation

We next explored the cellular origin of the subthreshold signal mixing. We repeated the patch-clamp experiment with a subset of the stimulation frequencies and a pharmacological blockade of synaptic input (see **Methods** for details). We found that blocking the synaptic ion channel currents attenuated, but did not abolish, the Δ*f* oscillation produced by the mixing of frequencies in the normal range of neural activity (i.e., *f*_1_ = 47 Hz and *f*_2_ = 57 Hz) (**Fig. 2A**). Synaptic blockade did not affect the mixing of frequencies beyond the normal neural range (i.e., *f*_1_ = 4.997 kHz and *f*_2_ = 5.007 kHz) (**Fig. 2B**). To exclude the possibility of a confounding contribution from presynaptic neurons driven above threshold at Δ*f*, we repeated the experiment with the same frequencies in the normal range of neural activity, but now with the stimulation currents applied intracellularly via the patch pipette to the recorded neurons. We found that the intracellular currents induced subthreshold membrane potential oscillations at their difference frequency (**Fig 2C**), confirming the single-cell origin of the subthreshold frequency mixing depolarization. To test whether the frequency mixing is linked to the nonlinearity of the voltage-gated ion channel conductance, we repeated the experiment with a pharmacological blockade of the voltage-gated sodium channels (using tetrodotoxin, TTX). We found that blocking TTX-sensitive sodium channels suppressed the mixing of membrane oscillations across the tested frequency range (**Fig. 2D-E**), implying their involvement in the subthreshold frequency mixing phenomenon (see **table S5** for all values and statistics for **fig. S2**).

**Fig. 2.**
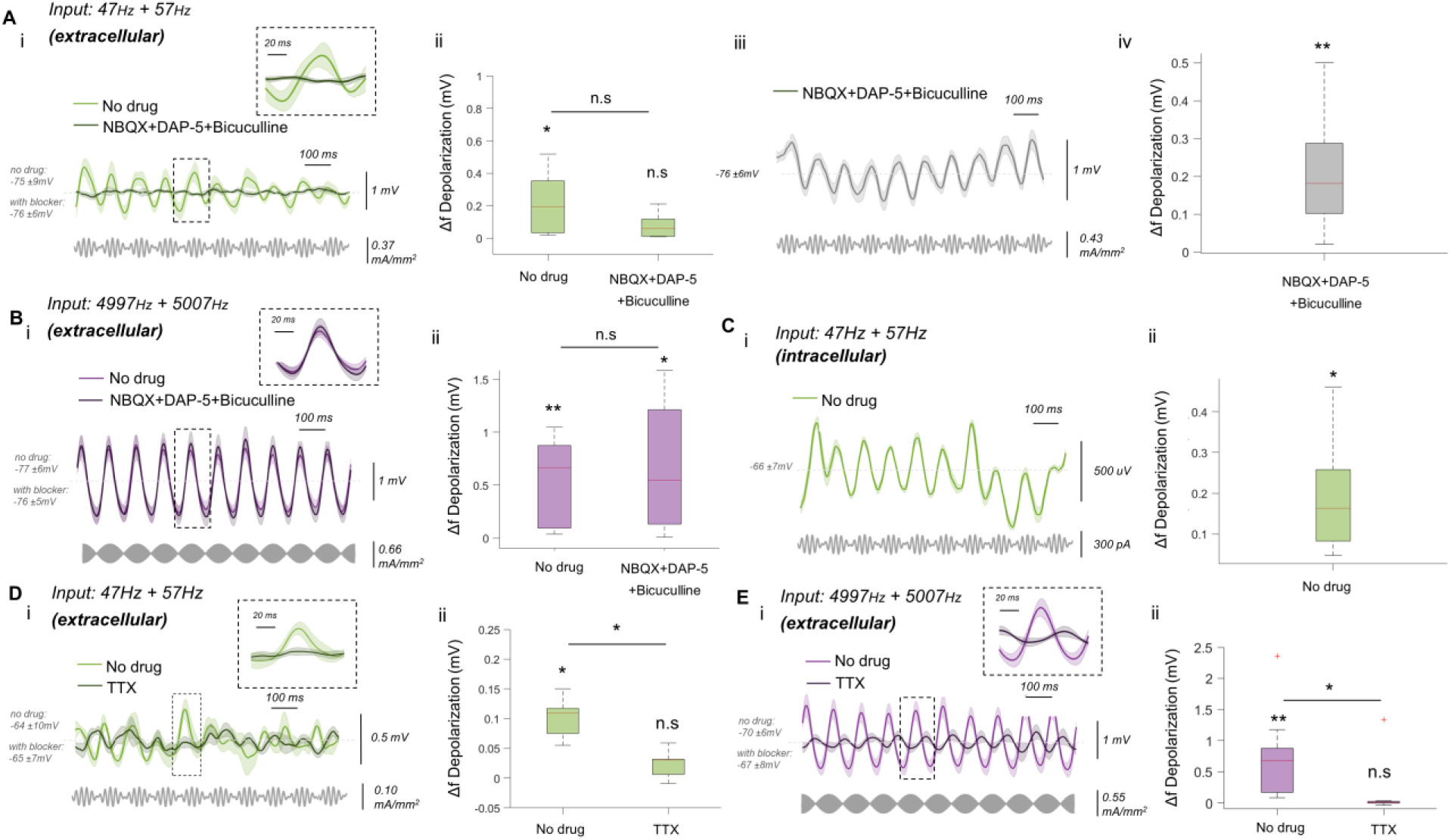
Origin of neuronal mixing characterized via pharmacological manipulation *ex vivo*. (**A**) (**i**) Membrane potential during extracellular electrical stimulation with *f*_1_=47Hz + *f*_2_=57Hz, before (color), and during (grey), pharmacological blockade of synaptic NMDA, AMPA, and GABA-A ion channel currents (shown are mean ± SEM). Raw membrane traces were filtered to remove stimulation artifact. Resting membrane potential mean ± SD is displayed. n=8 cells. Bottom: Applied stimulation current waveform. Zoom view of boxed region at the difference frequency (Δ*f*) is included. (**ii**) Boxplot of RMS amplitude of the induced neural oscillation at Δ*f* during the stimulation in (i) without drug and with drug. RMS values were baseline-subtracted. (**iii**-**iv**) same as (i-ii) but with higher current density stimulation. * above each box indicates significant oscillation at Δ*f* relative to measurement’s IMD at Δ*f*. *, p < 0.05, **, p < 0.005 two sample t-test. * between boxes indicates difference between drug conditions, paired t-test. n.s, non-significant. n(IMD/membrane potential)=8/8 recordings/cells. (**B**) As in (Ai-ii) but during stimulation with *f*_1_ = 4997Hz + *f*_2_ = 5007Hz. n=9 cells. n (IMD/membrane potential) = 8/9 recordings/cells. *, p < 0.05; **, p < 0.005. (**C**) As in (Ai-ii) but with intracellular electrical stimulation with *f*_1_=47Hz + *f*_2_=57Hz. (**D**) As in (Ai-ii) but before (color), and during (grey), pharmacological blockade of TTX-sensitive conductance. n(IMD/ membrane potential) = 7/7 recordings/cells. (**E**) As in (D) but during stimulation with *f*_1_ = 4997Hz + *f*_2_ = 5007Hz. n(IMD/ membrane potential) = 11/11 recordings/cells. Boxplots: central mark, median; box edges, 25th and 75th percentiles; whiskers, 1.5x interquartile range; ‘+’, datapoints outside this range.

### Mixing of endogenous membrane potentials in individual neurons

After confirming that neurons mix exogenous signals, we next examined whether they also mix endogenous (spontaneous) subthreshold membrane potential fluctuations. Subthreshold rhythmic fluctuations are common in neural cells (Buzsáki & Draguhn, 2004a; Giocomo et al., 2007; Hutcheon et al., 1996; Richardson et al., 2003). If the membrane potential depolarizes at frequencies *f*_1_ and *f*_2_ (*f*_2_>*f*_1_), and those frequencies are mixed by the membrane, then the instantaneous phases of the frequencies *f*_1_, *f*_2_ and Δ*f* = *f*_2_-*f*_1_ must be dependent, i.e., the frequency triplet must show a three-way, but not a pairwise, phase dependency (Haufler & Paré, 2019). Similarly, the instantaneous phases of the triplet *f*_1_, *f*_2_ and ∑*f* = *f*_1_+*f*_2_, the triplet *f*_1_, Δ*f*, and ∑*f*, and the triplet *f*_2_, Δ*f*, and ∑*f*, must also show a three-way phase dependency. Together, the four frequencies comprise a frequency mixing quadruplet. We repeated the *ex vivo* experiment but now we recorded the transmembrane potentials without electrical stimulation and then assessed the joint phase interaction of all possible frequency-mixing quadruplets (i.e., roots: *f*_1_, *f*_2_; products: Δ*f*, ∑*f*) for frequencies within the normal range of neural activity (i.e., up to 250 Hz) using a nonparametric test based on the Lancaster interaction measure (Lancaster, 2004; Rubenstein et al., 2016; Sejdinovic et al., 2013) (**Fig. 3A**; see **fig. S4** for sensitivity analysis). The statistical interaction test comprised a 3-way interaction measure that was compared against a generated null distribution, which upon rejection indicates that the joint probability distribution of the 3 frequency components cannot be factorized (either into pairwise or marginal distributions). To examine the mixing strength, we also defined a heuristic measure of the joint higher-order interaction (JHOI) strength based on the ratio of the interaction measure and the 95% quantile of the null distribution. See Methods for details of the nonparametric test.

**Fig. 3.**
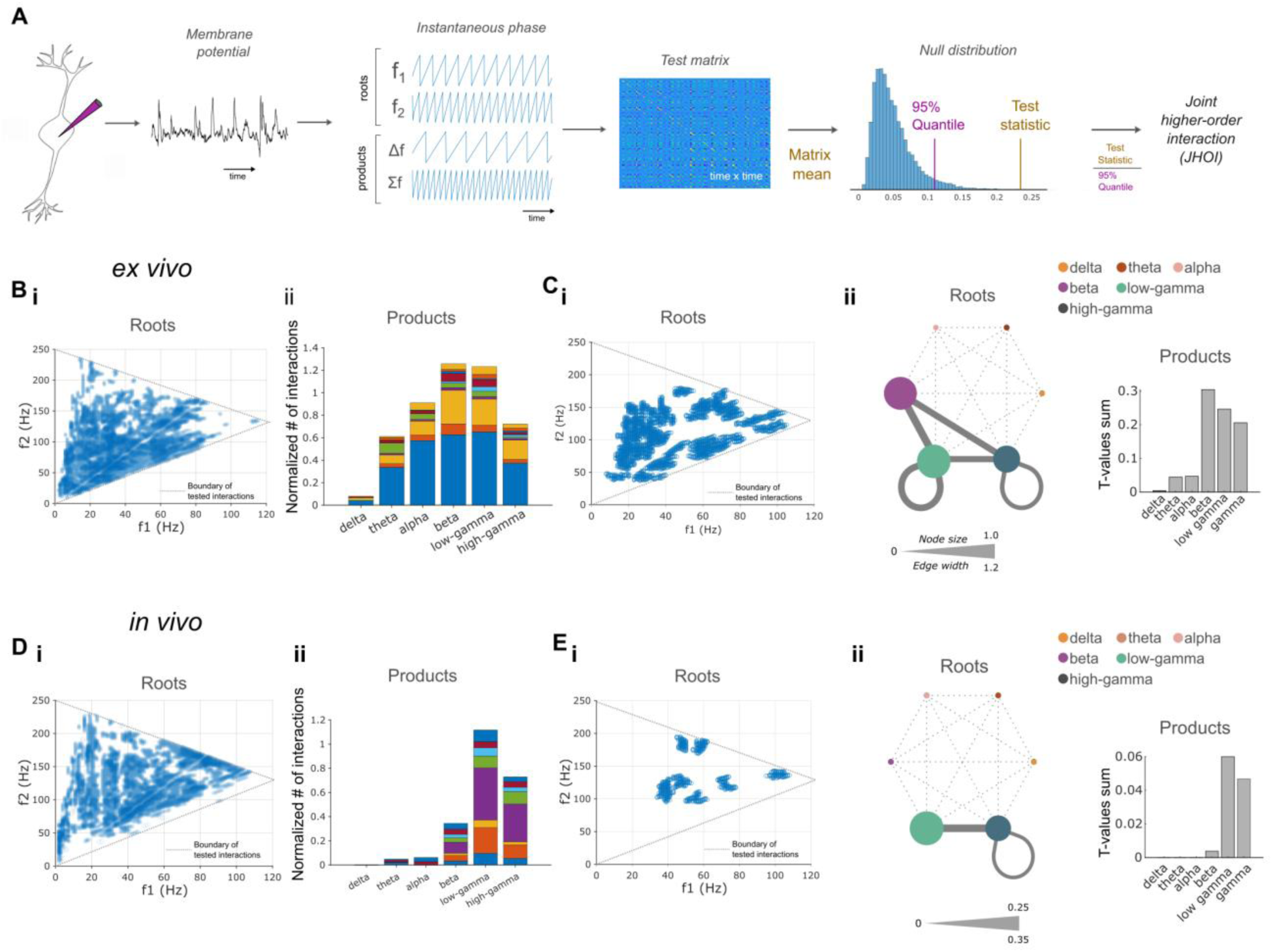
Endogenous membrane potential mixing in individual neurons *ex vivo* and *in vivo*. (**A**) Workflow to assess endogenous frequency mixing in cell membrane potential. For a given trace of endogenous membrane potential, the instantaneous phases of four frequency mixing components (roots: *f*_1_, *f*_2_ > *f*_1_, products: Δ*f* = *f*_2_ – *f*_1_, ∑*f* = *f*_1_ + *f*_2_) are extracted. Each subset of three phases (triplet) are used to construct a test matrix, which, when compared to a null distribution, defines the joint higher-order interaction (JHOI) strength heuristic. (**B**-**C**) *Ex vivo* recordings. (**B**) Frequency quadruplets with significant mixing in individual cells, assessed via permutation cluster test for each cell. Shown are (i) significant mixing root frequencies overlaid, (ii) the corresponding number of significant mixing interactions stacked by cell and normalized by the number of tested interactions per frequency band. n=20 cells. (**C**) Frequency quadruplets with significant mixing consistent across the cells, assessed via group-level permutation cluster test, showing (i) distribution of root frequencies overlaid, (ii) root frequencies stratified by frequency bands (network plot**^†^**), and distribution of frequency mixing products (bar chart). n=10 cells. (**D**-**E**) *In vivo* recordings. (**D**) As in (B) but *in vivo*. (**E**) As in (C) but *in vivo*. **^†^**Network plots: node size proportional to normalized sum of t-values of significant quadruplets within band (t-test vs. surrogate), edge width as in node size for roots shared between bands.

We found significant frequency mixing in the spontaneous transmembrane fluctuations of individual neurons spreading across a wide range of root frequency clusters (**Fig. 3Bi**, cluster permutation test computed for each cell individually, see Methods for details). The mixings produced a broad range of new frequencies peaking at the beta and low-gamma bands. **Fig. 3Bii** shows the number of significant mixing interactions stacked by cell and normalized by the number of tested interactions per frequency band. The mixing clusters within and between the beta and gamma bands were consistent across the cells (**Fig. 3C**, computed via a permutation cluster test at the group level against surrogate data), suggesting that certain frequencies are more commonly mixed. Moreover, by taking the average mixing across all mixing interactions up to 250 Hz, we found significantly stronger mixing in the neural membrane potentials vs. surrogate data (p=0.0027, paired t-test, n=10 cells). Adding a pharmacological blockade of the synaptic ion channel currents (as before) reduced the overall frequency mixing strength in the neural membrane potentials by approximately 30% (**fig. S5A**), particularly in a subset of high-frequency mixing clusters (**fig. S5B-C**).

To explore whether endogenous membrane potential mixing also occurs in the intact rodent brain, we recorded the membrane potentials of individual neural cells *in vivo*, using automated whole-cell patch-clamp recordings (Kodandaramaiah et al., 2012a) and deployed the same computation strategy to assess the phase dependency in all possible frequency-mixing quadruplets. We found significantly stronger frequency mixing in the spontaneous membrane potentials of the cells (mean JHOI amplitude across all mixing interactions up to 250 Hz 0.596 ± 0.168, mean ± st.d.; n = 8 cells from 8 animals; p = 0.009; compared to surrogate). The membrane potentials of individual cells exhibited a variety of frequency mixing clusters, as in brain slices. The mixings produced a narrower range of frequencies in the beta and gamma bands (**Fig. 3D**). Across the cells, the frequency mixing clusters were consistent in the gamma bands (**Fig. 3E**), supporting an earlier report using LFP recording (Haufler & Paré, 2019).

### Mixing of endogenous neural network oscillations in the human brain

After establishing the cellular origin of the neural frequency mixing phenomenon, we aimed to test whether the phenomenon exists in human brain oscillations, expanding on earlier evidence of mixing of endogenous neural network oscillations in rodents (Ahrens et al., 2002; Haufler & Paré, 2019). Neural oscillations are ubiquitous in the human brain (Buzsáki et al., 2013). We focused on the most salient human brain oscillation, i.e., the posterior alpha oscillation that can be readily observed in EEG during an awake eyes-closed state (Berger, 1929) and its established role in visual attention modulation (Carrasco, 2011; Fries et al., 2001). We recorded awake eyes-closed EEG in healthy human subjects (n = 20, mean age 29.3 ± 12.2 st.d., 6 females) and subsequently measured their visual attention control using a feature-matching task (Hampshire et al., 2012). We used the same computation strategy to examine the phase interaction in all possible frequency mixing quadruplets (up to 45 Hz) in EEG electrodes at a subset of sites in the parieto-occipital, temporal, and prefrontal regions (i.e., Pz, Oz, T7, T8, FP1, FP2 of the international 10-10 system) implicated in visual attention control (**Fig. 4A**).

**Fig. 4.**
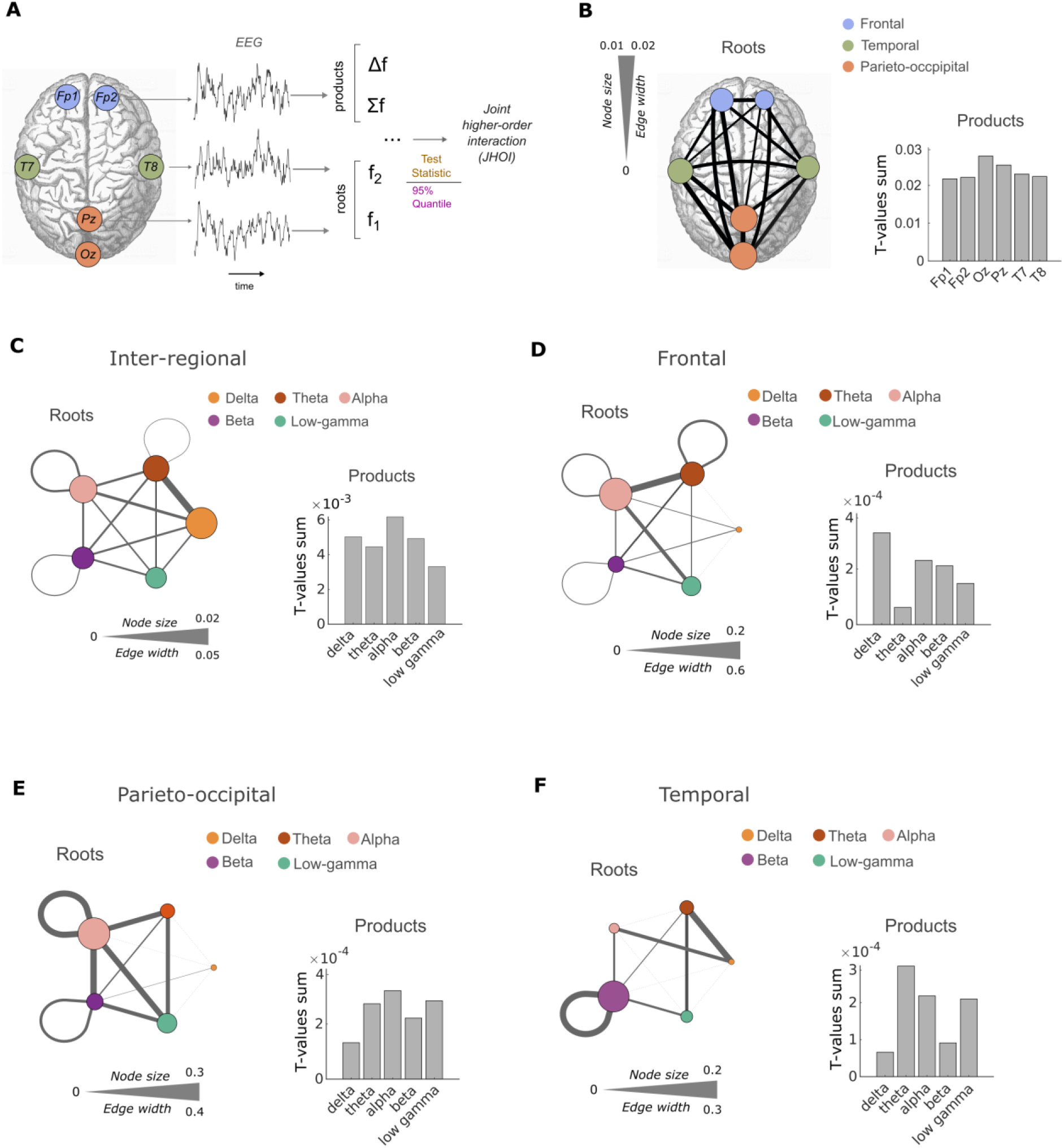
Mixing of endogenous neural network oscillations in the human brain EEG. (**A**) Illustration of workflow for assessing endogenous frequency mixing in the human brain EEG. Similar to (3A) but with instantaneous phases of the four frequency mixing components extracted from one or multiple cortical locations (in this example, roots: Pz and T8; products: Fp2). (**B**-**C**) Mixing of neural network oscillations between cortical sites. (**B**) Spatial topology of frequency mixing (significant at group-level against surrogate) for roots**^†^** (network plot) and products (bar chart). (**C**) Bands topology of frequency mixing (significant at group-level against surrogate) for roots**^†^** (network plot) and products (bar chart). (**D**-**F**) Mixing of neural network oscillations within cortical sites. (**D**) Bands topology of frequency mixing (significant at group-level against surrogate) in the frontal brain region for roots**^†^** and products (network plot). (**E**) As in (D) but in the parieto-occipital brain region. (**F**) As in (D), but in the temporal brain region. **^†^**Network plots: node size proportional to normalized sum of t-values of significant quadruplets within band/channel (t-test against surrogate), edge width same as node size but for roots shared between bands/channels. Boxplots: central line, median; circle, mean; whiskers, interquartile range; grey dots, outliers.

We found a robust mixing of neural network oscillations between sites of the human brain (JHOI amplitude 0.4 ± 0.008, mean ± st.d., p = 1.6e-16, paired t-test vs surrogate data). The spatial topology of the mixing is shown in **Figure 4B**, and the frequency band topology is shown in **Figure 4C**. The inter-site mixings occurred between all brain regions and frequency bands (**fig. S6A-C**) yet were stronger between the delta and theta bands. We also found mixing within brain sites, i.e., between local oscillations (JHOI amplitude 0.4 ± 0.02, mean ± st.d., p = 7.2e-12). The local mixings also occurred in all brain regions and frequency bands (**fig. S6D-F**), yet each brain region displayed a unique frequency band mixing pattern (**Fig. 4D-F**). The frontal region was dominated by theta-alpha mixing, parieto-occipital, alpha mixing, and temporal regions by beta mixing. The strength of oscillation mixing was not correlated with the oscillation power (**fig. S7A**), implying that mixing is a distinct feature of brain oscillation dynamics.

As expected, a strong alpha oscillation dominated the participants’ awake-eyes-closed EEG (**fig S7B**). We found that the mixing strength of this alpha oscillation was correlated with the participants’ visual attention capacity, indexed by the score in the subsequential feature-matching task (**Fig. 5A**, R^2^= 0.363, p=0.017, linear regression). A further investigation revealed that the alpha oscillation mixings associated with visual attention were specific to those with the beta oscillation (**Fig. 5B**, R^2^= 0.497, p=0.003, linear regression with Bonferroni correction for multiple comparisons). These alpha-beta mixings were strongest within the occipital cortex and between the occipital (alpha oscillation) and parietal (beta oscillation) cortices (**Fig. 5C**, repeated-measures ANOVA F(2.48), p=1.06e-5), producing new oscillations that were strongest in the gamma band (posteriorly) and weakest in the delta band (**Fig. 5D**, repeated-measures ANOVA Oz-Oz F(2.6), p=1.18e-5; Oz-Pz F(2.98), p=8.5e-7). These results suggest that the visual attention capacity may be modulated by frequency mixing strength, i.e., the efficiency by which the salient posterior alpha oscillation is mixed to augment local synchronization in the gamma band.

**Fig. 5.**
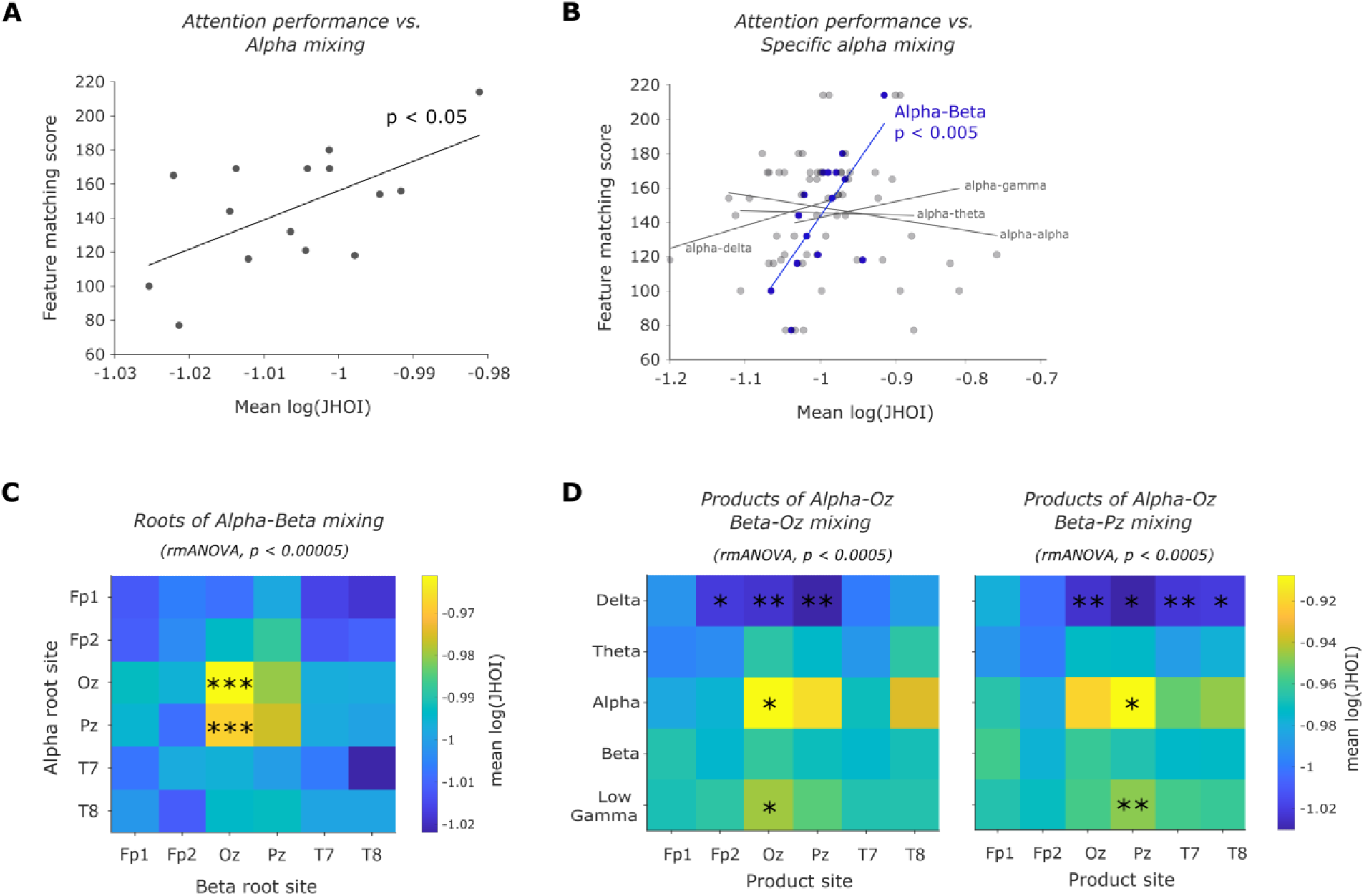
Human network oscillation mixing correlates with visual attention control. (**A**) Participants’ feature matching score vs strength of all alpha oscillation mixings (log JHOI averaged across all inter-site and local mixings), R^2^= 0.363, p=0.017, linear regression. (**B**) Participants’ feature matching score vs strength of alpha oscillation mixing with specific bands, showing significance at alpha-beta mixing, R^2^= 0.497, p=0.003, linear regression, Bonferroni corrected for multiple comparisons. (**C**) Topology of attentional correlated alpha-beta mixing roots, showing strongest mixing within Oz and between Oz alpha and Pz beta. ***, p<0.001, post-hoc paired t-test comparisons. (**D**) Topology of alpha-beta mixing products originated in Oz alpha-Oz beta (left panel) and Oz alpha-Pz beta (right panel), showing strongest products in posterior alpha and gamma bands and weakest products at delta band. *, p<0.05, **; p<0.01; post-hoc paired t-test comparisons. rmANOVA, repeated measures ANOVA.

## Discussion

The electrodynamics of the neural cell membrane underpins the brain’s computational functions. Neurons sum minuscule electrochemical synaptic inputs across their dendritic trees to yield integrated transmembrane potentials that, above a threshold, evoke action potentials and subsequent electrochemical axonal outputs (Aidley, 1978; Bean, 2007; Colbert & Pan, 2002). Nonlinear processes in the synapses and dendritic tree have been shown to enable complex signal processing (Beniaguev et al., 2021; Koch & Segev, 2000; Silver, 2010) while subthreshold rhythms functionally coordinate the processing across distributed neurons (Buzsáki & Draguhn, 2004a; Giocomo et al., 2007; Hutcheon et al., 1996; Richardson et al., 2003). In this paper, we report that electrodynamics of the neural cell membrane involves mixing subthreshold rhythms, thereby actively producing new frequencies.

Our report expands on previous reports of frequency mixing in rodents (Ahrens et al., 2002; Haufler & Paré, 2019) by demonstrating the single cell origin of the phenomenon and its existence and functional relevance in humans. Our report strengthens hitherto evidence of single cell multiplication based on the coincidence detection capacity (Gabbiani et al., 2002; Groschner et al., 2022; Lavzin et al., 2012; Poleg-Polsky and Diamond, 2016) by demonstrating the frequency mixing capacity. Our results suggest that the mixing of subthreshold frequencies occurs in the currents of the voltage-gated ion channels, and that the mixing process is highly efficient. Specifically, the threshold for inducing spike trains by mixing frequencies in the normal neural range was similar to that by direct stimulation at the target frequency. The mixing efficiency of kHz electrical stimulation was 1.5-2 times weaker than with frequencies within the normal range. These results help elucidate the mechanism by which temporal interference of kHz electric fields stimulates neural activity (Grossman et al., 2017).

Neural oscillations are ubiquitous in the human brain (Buzsáki et al., 2013) and are implicated in regulating behavioral states (Doiron et al., 2016), coordination of multisensory processing (Keil & Senkowski, 2018), and cognitive processes, such as memory and consciousness (Ward, 2003). Aberrant oscillations have been associated with almost all neurological and psychiatric disorders (Başar, 2013; Nimmrich et al., 2015; Uhlhaas & Singer, 2006; Vanneste et al., 2018). The frequencies of neural oscillations have been thought to emerge from competition between local oscillators since different oscillations can naturally emerge in neural networks with different cell-type compositions (Buzsáki & Draguhn, 2004b). (Buzsáki & Draguhn, 2004b). Our findings, together with the original studies in rodents (Ahrens et al., 2002; Haufler & Paré, 2019), suggest that individual neurons can control the frequencies of their network oscillations via a membrane-mixing phenomenon.

The brain has traditionally been modelled as a complex system composed of pairwise interactions among different elements. There is a growing acknowledgment that such a theory offers a limited description of brain function (Battiston et al., 2021; Lambiotte et al., 2019). High-order interactions, which involve groups of three or more elements, are increasingly recognized as foundational to the architecture of many complex systems and appear to play a pivotal role in cognition (Gatica et al., 2021; Tononi et al., 2016). Here, we posit that frequency mixing may be a fundamental mechanism for integrating high order information across spatiotemporal scales. Our results suggest that the topology and strength of frequency mixing in the human brain has behavioral relevance We show that the mixing of the salient posterior alpha-beta oscillations to produce gamma-band oscillations correlates with the visual attention state. There is substantial evidence that posterior alpha oscillation modulates visual cortex excitability and, consequentially, visual perception and attention (see, for example (Sadaghiani & Kleinschmidt, 2016)). The mixing of alpha and beta oscillations to produce gamma oscillations, which have been linked to spiking activity, may underpin the mechanism of excitability modulation. Our data show that each brain region has a unique pattern of mixing oscillations modulated by inter-regional mixing, suggesting a mechanism coupling local and global oscillations (see also (Doiron et al., 2016) image-reject mixing in communication theory). In this study, we did not directly test the link between frequency mixing in the EEG oscillations and the frequency mixing in the subthreshold membrane potential of individual neurons. The fact that extracellular signals such as LFP and EEG oscillations originate predominantly from synchronous subthreshold activity of individual neurons (Buzsáki et al., 2012) allows a conceptional link between the human EEG and single neuron patch clamp results. However, future studies using concurrent single-cell and network-level recording should further elucidate this link and could shed light on the precise mechanisms underpinning integration of high-order computations across different scales.

The functional role of neural oscillations has been linked to the coordination of spiking activity between brain sites because task-induced synchronization, i.e., phase alignment, also known as functional connectivity, has been observed (Ward, 2003). Our results imply that individual neurons could directly utilize these oscillations to perform advanced computational operations such as phase detection and (de)multiplexing that, until now, have only been seen in modern telecommunication.

## Materials and Methods

### Single-cell investigation *ex vivo*

#### Animals

All mice were male C57BL/6, aged between 4-12 weeks. Mice were housed in standard cages in the Imperial College London animal facility, with ad libitum food and water in a controlled light-dark cycle environment, with standard monitoring by veterinary staff. The Imperial College London’s Animal Welfare and Ethical Review Board approved all animal procedures, and all experiments were performed in accordance with relevant regulations/according to the United Kingdom Animals (Scientific Procedures) Act 1986.

#### Brain slice preparation

Anaesthesia of mice was achieved via intraperitoneal injection of 100 mg/kg ketamine and 10 mg/kg xylazine. Mice were transcardially perfused with 0–5 degree C carbogenated dissection artificial cerebrospinal fluid (aCSF) containing (in mM): 108 C5H14ClNO, 3 KCl, 26 NaHCO3, 1.25 NaH2PO4, 25 Dextrose, 3 C3H3NaO, 1 Mgcl2, and 2 CaCl2. 350-μm-thick coronal slices around the somatosensory cortex were prepared using a Vibratome (Campden Instruments LTD, Loughborough, UK). Following sectioning, slices recovered for 2-4 hours at room temperature in carbogenated bath aCSF containing (in mM): 120 NaCl, 3 KCl, 23 NaHCO3, 1.25 NaH2PO4, 1 Mgcl2, and 2 CaCl2. Following recovery, slices were placed in the recording chamber of an upright microscope (Scientifica, Uckfield, UK), and held down using a harp (Multi Channel Systems, Reutlingen, Germany). The recording chamber was continually perfused with room temperature carbogenated bath aCSF throughout the experiment.

#### Whole-cell patch clamp recording

Patch electrodes were pulled from filamented thin-walled borosilicate glass capillaries (World Precision Instruments, Hitchin, UK) using a horizontal Flaming–Brown micropipette puller (P1000; Sutter Instruments, Novato, CA, USA). Electrode tip resistance immediately after pulling ranged from 5-8 MΩ. Whole cell recordings were taken in a current clamp mode (no holding current) using a patch clamp amplifier (MultiClamp 700B; Molecular Devices Ltd). The recorded traces were digitalized using a digitiser (Digidata 1550b; Molecular Devices Ltd) at 100 kS/s rate. Recordings were obtained from the soma of L2/3 cortical neurons in coronal brain slices. Patch electrodes were filled with internal solution containing (in mM): 130 KMeSO4, 8 NaCl, 2 KH2PO4,2 Dextrose and 10 HEPES. Following successful break-in and before each electrical stimulation, a protocol comprising of current steps increasing in amplitude by 50pA was run, to determine viability of the cell, resting membrane potential, and threshold for AP firing.

#### Electrical stimulation

Stimulating current waveforms were generated using a custom-written MATLAB GUI via an arbitrary voltage waveform generator (USB-6216 BNC; National Instruments, Newbury, UK) sampled at 250 kS/s and an isolated constant current source (LCI1107; Soterix Medical Inc, New York, NY, USA). The current waveforms were applied to the tissue via a 0.25mm diameter platinum wire electrode (VWR, Lutterworth, UK) positioned 50-100μm from the recorded neuron, touching the slice, on L2/3 cortex. The stimulating electrode was paired with two remote 2x2mm Ag/AgCl electrodes (VWR, Lutterworth, UK) in a y-shape configuration. The stimulation protocol included a series of two symmetrical biphasic sinusoidal waveforms with the same amplitude and a difference frequency (typically 10Hz). The two sinusoidal waveforms were summed before they were applied to the tissue resulting in a combined waveform that oscillates at the mean frequency and has an envelope amplitude that changes periodically at the difference frequency. We tested stimulation conditions with an average (“mean”) frequency between 10Hz and 5,000Hz and a range of current amplitudes (0.5mA current steps). Each stimulation condition was applied for 5s with 0.5s ramp-up, 0.5s ramp-down, and a 5s stimulation-free period between consecutive stimulations (baseline). The order of the stimulation conditions was randomized between recordings. Current density at the electrodes was estimated by dividing the applied current amplitude by the cross-sectional area of the stimulation electrodes.

#### Pharmacological manipulation

Blockade of synaptic ligand-gated ion channels was achieved by application of an AMPA receptor antagonist (NBQX; Merck Life Science UK Ltd, #N183), an NMDA receptor antagonist (DAP-5; Merck Life Science UK Ltd, #A8054) and GABAA antagonist (Bicuculline; Merck Life Science UK Ltd, #14340) to the bath aCSF being perfused to the slice. The resulting bath aCSF solution contained (in μM): 10 NBQX, 50 DAP-5, and 10 Bicuculline. One minute of spontaneous membrane potential activity was recorded prior to drug application and 10 minutes following, to observe elimination of postsynaptic potentials. Elimination of postsynaptic potentials was observed in 100% of cells included in analysis. Blockade of voltage-gated sodium channels was achieved by application of Tetrodotoxin (TTX) (Abcam Ltd, #ab120054) to the bath aCSF being perfused to the slice. The resulting aCSF solution contained in μM): 1 TTX. Membrane potential response to current steps of 50pA was recorded prior to drug application and 10 minutes following, to observe elimination of action potentials (Aps). Disappearance of APs was observed in 100% of cells included in analysis. Post drug stimulation and recordings commenced 10 minutes following drug application in all cases.

### Single-cell investigation *in vivo*

#### Animals

All mice were male C57BL/6, aged between 4-12 weeks. Mice were housed in standard cages in the Massachusetts Institute of Technology (MIT) animal facility, with ad libitum food and water in a controlled light-dark cycle environment, with standard monitoring by veterinary staff. All animal procedures were approved by the MIT Committee on Animal Care (CAC, Protocol Number: 1115-111-18), and all experiments conformed to the relevant regulatory standards.

#### Surgery

On the day of the experiment, the mice were injected with Meloxicam (1mg/kg) and buprenorphine (0.1mg/kg) and anesthetized with 1%–2% (vol/vol) isoflurane in oxygen. Ophthalmic ointment (Puralube Vet Ointment, Dechra, KS; USA) was applied to the eyes. The scalp and was shaved and sterilized with Betadine and 70% ethanol, a custom-made head-plate was attached using dental cement (C&B Metabond, Parkell, NY; USA), and a craniotomy was performed.

#### Whole-cell patch clamp recording

*In vivo* whole cell patching in current clamp mode was conducted in the cortex (depth of ∼500 μm below the dura) of anesthetized mice with an autopatcher (Kodandaramaiah et al., 2012b). Whole cell recordings were taken in a current clamp mode (no holding current) using a patch clamp amplifier (MultiClamp 700B; Molecular Devices Ltd, Wokingham, UK). The recorded traces were digitalized using a digitiser (Digidata 1550b; Molecular Devices Ltd, Wokingham, UK) at 25kS/s rate. Patch electrodes were pulled from thin-walled borosilicate glass capillary tubing using a pipette puller (P97; Sutter Instruments, Novato, CA, USA). Tip electrode resistance was 4.6–7.4 MΩ in artificial cerebrospinal fluid (ACSF), containing (in mM): 126 NaCl, 3 KCl, 1.25 NaH2PO4, 2 CaCl2, 2 MgSO4, 24 NaHCO3 and 10 glucose. The patch electrode internal solution consisted of (in mM) 122.5 potassium gluconate, 12.5 KCl, 10 KOH-HEPES, 0.2 KOH-EGTA, 2 Mg-ATP, 0.3 Na3-GTP, 8 NaCl (pH 7.35, mOsm 296. Following successful break-in and before each electrical stimulation, a membrane test was run on the cell to gather membrane statistics. Furthermore, a protocol comprising of current steps increasing in amplitude by 50pA was run, to determine viability of the cell.

### Human investigation

#### Participants

Twenty healthy adults (14 males, 6 females; mean age 29.29, range 18-70 years) were included. Written informed consent was obtained for all participants judged to have capacity according to the declaration of Helsinki. Human data was collected under approval by the West London Research Ethics Committee (09/HO707/82).

#### EEG recording

A 32-channel active electrode standard actiCAP (Easycap) was used to acquire 5 minutes of eyes-closed resting state EEG data. Measurements were taken from nasion to inion and from left to right tragus to position the cap in accordance with the international 10-20 system with the vertex electrode (Cz) in the centre. The cap was secured with a chin strap to maintain positioning. A small amount of conductive electrode gel was applied to each electrode using a blunt needle. Impedances for ground (Fpz) and reference (Fz) electrodes were maintained at <5kΩ and aimed to be kept under 50kΩ across all other electrodes. Signals from the electrodes were amplified using the actiCHamp system and data were recorded using the BrainVision Recorder software (Brain Products GmbH, Gilching, Germany) at a sampling rate of 1 kHz. Six channels were exported for further analysis as representative of the frontal (Fp1, Fp2), temporal (T7, T8) and parieto-occipital (Pz, Oz) regions. One minute of artifact-free resting-state EEG for each subject was then analysed using the methodology introduced below. For spectral analysis, the traditional EEG bands were defined as follows: delta (0-4Hz), theta (4-8Hz), alpha (8-13Hz), beta (13-30Hz) and low-gamma (30-45Hz).

#### Visual attentional control task

Visual attentional control was assessed using a feature-matching task delivered on a touchscreen tablet device using a custom-programmed application. The task included two grids presented side by side each containing a series of shapes. The shapes presented in each grid were identical for half of all trials and differed by one shape in the other half of trials. The participant requested to state whether the shapes match or mismatch. The first trial contained one shape in each grid. The number of shapes increased with each correct response and decreased with each incorrect response. The task lasted for 90 seconds during which the participants solved as many trials as possible. The main outcome measure was the total score. Population mean = 131.35, SD = 32.79 (Hampshire et al., 2012).

### Analysis

#### Exogenous frequency mixing

All data were analyzed in Matlab (Matlab 2019a, The MathWorks Inc.).

#### Characterization of measurement’s intermodulation distortions (IMDs)

Nonlinearity in the stimulation hardware can result in frequency mixing that can confound the measured neural signals at the frequency mixing products (i.e., Δ*f*, ∑*f*, 2*f*_1_, 2*f*_2_, and 2Δ*f*), aka intermodulation distortions (IMDs). Additional IMDs can arise from the recording hardware due to the large stimulation voltage at the amplifier input that risks shifting the dynamic range to a nonlinear gain. In our case, IMDs from the recording hardware were neglected as the amplifier operated in its linear gain range (tested by applying amplitude-modulated waveforms at different voltages directly to the amplifier). Whereas a high-pass filter could mitigate artifacts in the difference frequencies at the output of the current sources, artifacts in the sum frequencies were more challenging to mitigate in our experiment due to the spectral proximity to the applied frequencies and the smaller signal-to-noise-ratio (the amplitude of the transmembrane potential drops with frequency (Deans et al., 2007)). We characterized the measurement IMDs by repeating the experiments without brain slices and computing the root mean square (RMS) amplitude of the frequency mixing product of interest as in ‘Subthreshold depolarization analysis’ (see below). The artifactual stimulation potentials recorded at the applied frequencies (*f*_1_, *f*_2_) were amplitude matched and had the same stimulation artifact amplitude as those measured in the *ex vivo* condition (p > 0.05, Wilcoxon rank sum test or two-sample t-test).

#### Subthreshold depolarization analysis

Induced subthreshold membrane oscillation amplitude at the frequency mixing products (i.e., Δ*f*, ∑*f*, 2*f*_1_, 2*f*_2_, 2Δ*f*, 2*f*_1_-*f*_2_, ∑*f*-2Δ*f*) was computed. The membrane potential trace in response to the maximum stimulation amplitude that had no APs was taken. The signals were filtered using a high-pass filter (Butterworth; 1Hz, order 3) and a low-pass filter (Butterworth; 50Hz, order 5) to remove DC shift and stimulation artifact. Traces were then bandpass filtered around the frequency of interest (±1Hz) (Butterworth; bandwidth 2Hz, order: 5). The average stimulation-induced subthreshold oscillation amplitude for each neuron was computed by computing RMS of the middle 1s of the filtered signal. For all analysis of subthreshold neural activity in the frequency and time domains, only recordings during stimulation where the neuron did not exhibit a suprathreshold response were included, to ensure that the values being obtained were representative of purely subthreshold neural polarisation for that neuron. The RMS amplitudes at the frequencies of interest were tested against artifactual amplitude from the measurement’s IMDs, and baseline amplitude at those frequencies (i.e., prior to the stimulation onset). Specifically, the baseline RMS amplitude was subtracted from the stimulation RMS amplitude and then statistically compared to the computed IMD RMS amplitude (also baseline subtracted) using the Wilcoxon rank sum test or two-sample t-test. See ‘Characterization of measurement’s intermodulation distortions (IMDs)’ for details about the IMD measurement.

#### Threshold of inducing action potential (AP) train at Δ*f*

To remove the stimulation voltage artifact at the applied frequencies, the traces were low pass filtered if the applied mean frequency was ≥100Hz (Butterworth: cut-off frequency: 25Hz; order: 5), or high pass filtered if <100Hz (Butterworth: cut-off frequency: 100Hz; order: 3). Threshold was defined as the lowest applied current density required to evoke an action potential (AP) train with at least three spikes significantly phase locked to the stimulation (tested using the Rayleigh test). APs were detected via a custom-made script using a peak finding algorithm. Thresholds were normalized to the threshold for direct stimulation with the difference frequency (10Hz) for each neuron.

#### Plotted traces

For plotting of mean membrane potential, individual membrane potential traces were high-pass filtered to remove offset (Butterworth: cut-off frequency: 1Hz; order: 3), and low pass filtered to remove stimulation artifact (Butterworth: cut-off frequency: 25Hz; order: 5). The baseline membrane potential prior to each stimulation was computed as the mean membrane potential during the 0.5s preceding the stimulation (i.e., during the inter-stimulation interval).

#### Endogenous frequency mixing

All data were analyzed in Matlab (Matlab 2021a, The MathWorks Inc.).

#### Overview

Frequency mixing, at the most basic level, involves the non-linear interaction of two root frequencies generating two product frequencies (the sum and difference of the root frequencies). The four frequencies form a frequency-mixing quadruplet and any subset of three frequencies within the quadruplet form a frequency-mixing triplet. The frequency-mixing triplets display a 3-way phase relationship where there exist no underlying pairwise phase relationships (Haufler & Paré, 2019), i.e., the joint distribution of three instantaneous phase signals cannot be factorised. However, given that a 3-way phase relationship can exist, a 4-way phase relationship cannot be directly detected since it would be confounded by any potential underlying 3-way phase relationships (the joint distribution of four instantaneous phase signals can be factorised). Thus, to detect frequency mixing, we (1) detected frequency-mixing triplets using the Lancaster interaction measure and Wild Bootstrap, (2) computed a heuristic measure of joint higher order interaction (JHOI) and (3) inferred the presence of frequency-mixing quadruplets using the 4 subset triplets that constitute a quadruplet. Note, quadruplets, but not triplets alone, can reveal the underlying root frequencies.

#### Procedure

We computed the test statistic of the joint interaction for each frequency triplet by first computing the Gram matrices of the three instantaneous phase traces (computed via a Morlet wavelet decomposition with a width of 15 as in (Haufler & Paré, 2019)) by embedding them into a reproducing kernel Hilbert space (RKHS) with a Gaussian kernel (Berlinet & Thomas-Agnan, 2004). We empirically centred the Gram matrices by subtracting the row and column averages and adding the mean of the matrix elements. We then computed the test matrix by an element-wise multiplication of the three matrices (i.e., a Hadamard product) and averaging the matrix elements. To heuristically estimate the strength of the test statistic of the joint interaction, we normalized the test statistic value by the 95% confidence interval of the null distribution (i.e., the distribution of the test statistic values under the null hypothesis), computed using a Wild bootstrap permutation (Mammen, 2007) to preserve the temporal dependency in the instantaneous phase data (we used at least 10,000 permutations, which we found to be stable in robustness tests). The resulting test statistic value approximates the joint high-order interaction (JHOI) between the three instantaneous phase traces, i.e., a higher JHOI corresponds to a stronger interaction between the phase traces.

To identify frequency quadruplet clusters with a group-level significance, we used a cluster permutation t-test. We computed the t-value of each frequency quadruplet’s JHOI (taking the median JHOI of each quadruplet’s four frequency triplets) across the recordings using a pair-wise t-test against a null distribution generated from surrogate data. The surrogate data was generated from the recording time series by resetting the phase delay of its frequency components in the Fourier space. We then clustered frequency quadruplets with a p-value threshold of 0.01 against a null distribution of cluster t-value sum, generated by permuting the condition’s label. See **fig. S4** for a characterization of the JHOI computation strategy using synthetic data.

#### Synthetic data

To generate oscillatory signals, we followed the methodology described by Haufler et al (Haufler & Paré, 2019). Briefly, an interval was defined for the frequency and amplitude of each signal (normally +/- 1Hz for a given frequency and 0.5-1 amplitude for a given signal). Using these intervals, a frequency vector and amplitude vector was generated using cubic spline interpolation at 0.8ms steps (negative values of frequency or amplitude were set to zero). Next, a phase variable was integrated over the signal, advanced at a rate proportional to the instantaneous frequency. The resulting phase was parsed to a cosine function to generate an oscillatory signal which is then scaled at each point in time by the instantaneous amplitude vector. Two oscillatory signals components were combined by their simple addition and via a non-linear activation function (here we used a quadratic function), *F*(*s*) = *B* + *Cs* + *Ds*^2^, where *s* = *S*_1_ + *S*_2_ is comprised of two oscillatory signals and expansion of this result gives, *F*(*S*_1_ + *S*_2_) = *B* + *CS*_1_ + *CS*_2_ + *DS*_2_^2^ + *DS*^2^ + 2*DS*_1_ *S*_2_. Finally, white noise *σ* is added to the resulting signal.

#### Surrogate Data

To generate surrogate data, we used the original time-series signal and shuffled the phase components (Schreiber & Schmitz, 2000). We first performed a fast Fourier transform (Matlab, fft function), take the absolute magnitude of the spectral density (Matlab, abs function), and finally perform an inverse fast Fourier transform (Matlab, ifft function). The surrogates retained (i) the spectral power distribution of each respective time-series but removed the phase relationships between frequency components and (ii) retained the temporal dependence necessary for comparative time-series. Note that this should not be confused with the generation of null distributions for the non-parametric test for Lancaster interaction.

#### Cluster permutation t-test

The cluster-based permutation t-test was used to identify significant frequency-mixing quadruplets at the group-level using their JHOI values. We computed an uncorrected t-test (paired or two-sample t-tests depending on data) for each quadruplet. We retained quadruples with p-value <0.01 and clustered them according to their similarity in terms of both their frequency difference and spatial distance. The similarity in frequency was defined such that for two quadruplets *a* and *b*, 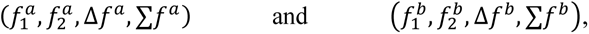 if their distance 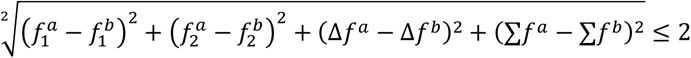 the quadruplets were considered neighbours. The similarity in space was defined by the EEG electrode locations, such that two quadruplets were considered neighbours if at least three frequencies from one quadruplet were recorded from the same electrodes as the corresponding frequencies in the second quadruplet. For example, *f*_1_*^a^* and *f*_1_*^b^* recorded from the same electrode, *f*_2_*^a^* and *f*_2_*^b^* recorded from the same electrode, and ∑*f^a^* and ∑*f^b^* recorded from the same electrode. For each cluster, the t-values are summed and then compared against a null distribution. The null distribution was generated by permuting the condition label 50,000 times and then re-clustering and calculating the maximum cluster t-value sum for each permutation. The t-sum of each cluster in the original data was then contrasted with the null distribution and significant (p<0.05) clusters are identified.

#### Statistical analysis

Values in text and supplementary tables are mean ± standard deviation (st.d). Statistical tests are specified in text/figure legends/supplementary tables. All t-tests are two tailed unless stated. In all plots, the * notation represents a statistical significance that survived Bonferroni correction if required.

## Acknowledgments

We thank Lok Fan for improving computational efficiency, Yuval Gal Shohet for discussions on JHOI measure, Dr Ho-Jun Suk for assisting with *in vivo* patch-clamp experiments, and Dr Carola Radulescu and Dr Sam Barnes for help in setting up the *ex vivo* patch-clamp experimental rig.

## Funding

Deutsche Forschungsgemeinschaft, DFG, German Research Foundation, Project-ID 424778381-TRR 295 (RP)

EPSRC award EP/N014529/1 supporting the EPSRC Centre for Mathematics of Precision Healthcare at Imperial College London (RP, FL, MB)

UK Dementia Research Institute—an initiative funded by the Medical Research Council (NG, CL)

Engineering and Physical Sciences Research Council, UK, EP/W004844/1 (NG)

National Institute for Health and Care Research, Imperial Biomedical Research Centre (NG)

## Author contributions

Conceptualization: NG, CL, RP

Methodology: CL, RP, FL

Investigation: CL, RP, EJM

Visualization: CL, RP, ER

Supervision: NG, MB, ESB, DJS

Writing: NG, CL, RP, ER, EJM

## Data and materials availability

All data, code, and materials used in the analysis are available from the corresponding author upon request.

(We will also deposit all the data and code on a public repository once the manuscript is in its final form, i.e., upon completion of the review and before publication)

---

**Fig. S1.**
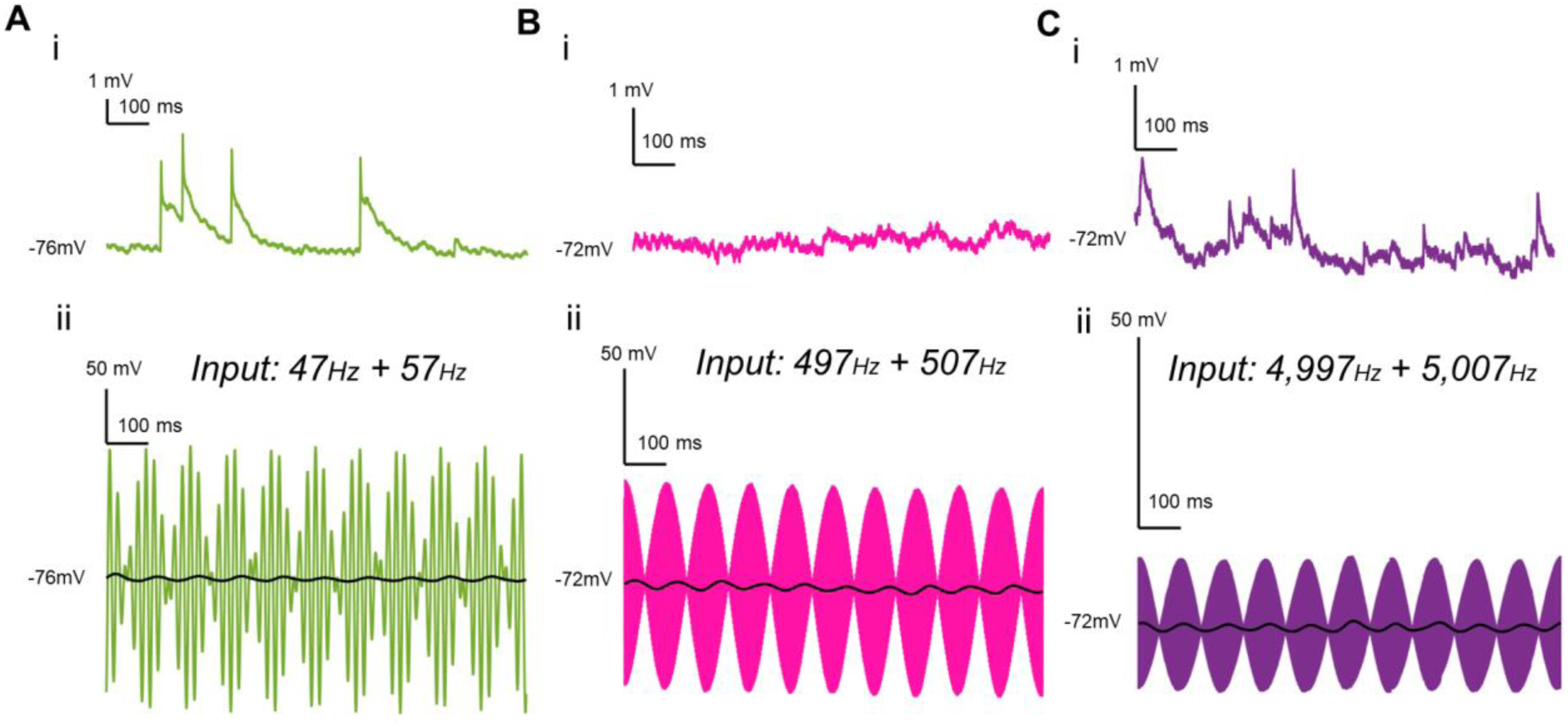
Representative membrane potential recordings during baseline (no stimulation) and stimulation. (A) Unfiltered recorded signal during (i) baseline (no stimulation) and (ii) stimulation (with low pass filtered recording overlaid (black)), for a representative neuron during stimulation with sinusoidal electrical currents *f*_1_=47Hz + *f*_2_=57Hz. Unfiltered stimulation recording is dominated by stimulation artifact. (B) As in (A) but for stimulation with *f*_1_=497Hz + *f*_2_=507Hz. (C) As in (A) but for stimulation with *f*_1_=4997Hz + *f*_2_=5007Hz. Recordings from different representative neurons are shown for each stimulation frequency.

**Fig. S2.**
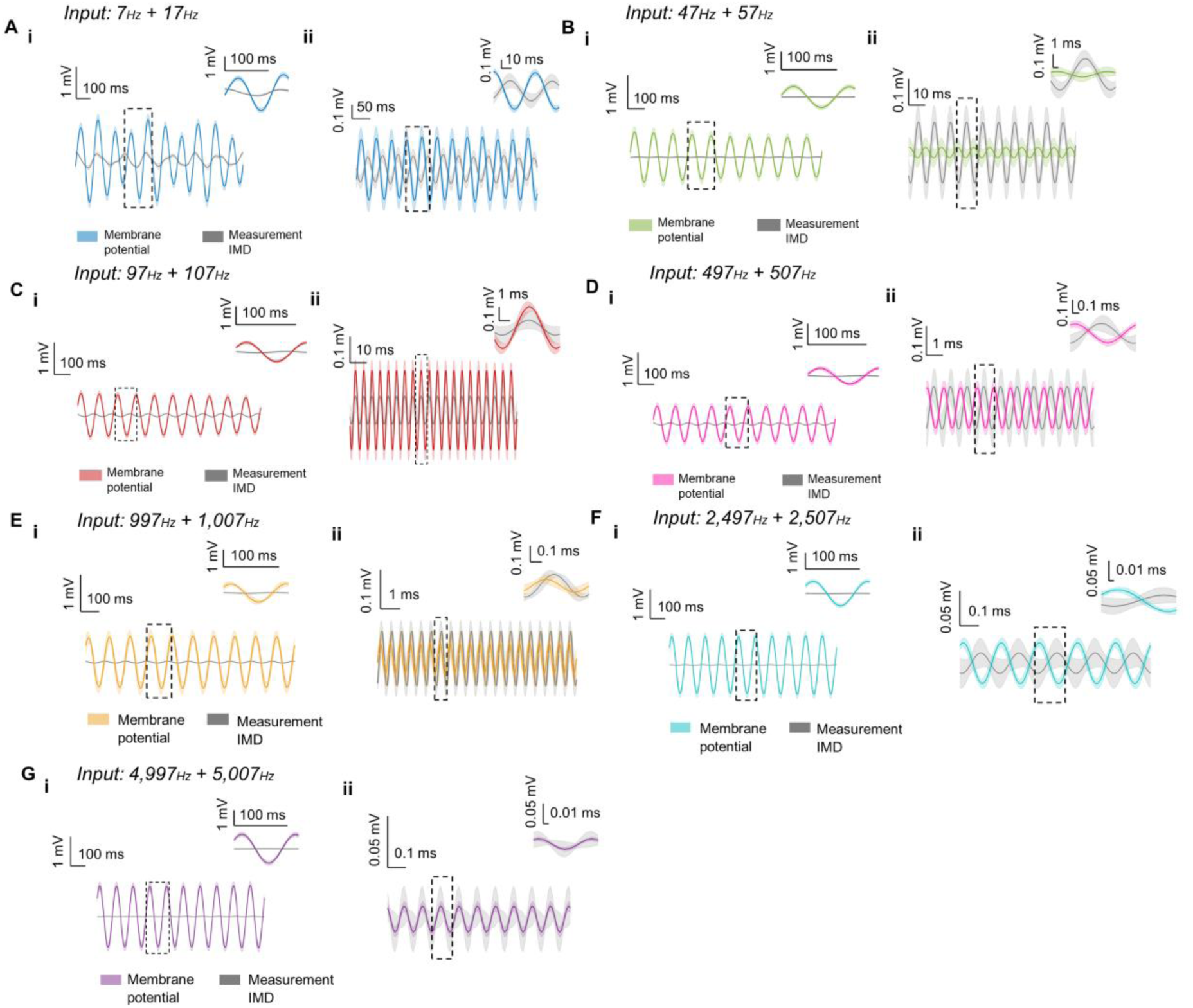
Mixing of exogenous membrane potentials in individual neurons *ex vivo* – membrane potential traces. (**A**) (i) Recorded signal at Δ*f* from neural cells (color) and without cells showing the measurement’s intermodulation distortion (IMD) at Δ*f* (grey) during stimulation with sinusoidal electrical currents *f*_1_=7Hz + *f*_2_=17Hz. Data shown are mean ± SEM. Individual membrane traces were filtered to remove stimulation artifact. Zoom view showing plot inside dashed box. (ii) As in (i), but data were filtered to isolate oscillation at the sum frequency (*∑f*). (**B**) As in (A) but for stimulation with *f*_1_=47Hz + *f*_2_=57Hz. (**C**) As in (A) but for stimulation with *f*_1_=97Hz + *f*_2_=107Hz. (**D**) As in (A) but for stimulation with *f*_1_=497Hz + *f*_2_=507Hz. (**E**) As in (A) but for stimulation with *f*_1_=997Hz + *f*_2_=1,007Hz. (**F**) As in (A) but for stimulation with *f*_1_=2,497Hz + *f*_2_=2,507Hz. (**G**) As in (A) but for stimulation with *f*_1_=4,997Hz + *f*_2_=5,007Hz. n (IMD/Membrane potential) = 20/27 (7/17Hz); 20/21 (47/57Hz); 25/27 (97/107Hz); 26/27 (497/507Hz); 21/25 (997/1,007Hz); 29/29 (2,497/2,507Hz); 28/29 (4,997/5,007Hz) recordings/cells. Current densities: 0.41±0.31 (7/17Hz); 0.30±0.24 (47/57Hz); 0.34±0.27 (97/107Hz); 0.34±0.22 (497/507Hz); 0.33±0.26 (997/1,007Hz); 0.45±0.34 (2,497/2,507Hz); 0.69±0.37 (4,997/5,007Hz) mA/mm^2^.

**Fig. S3.**
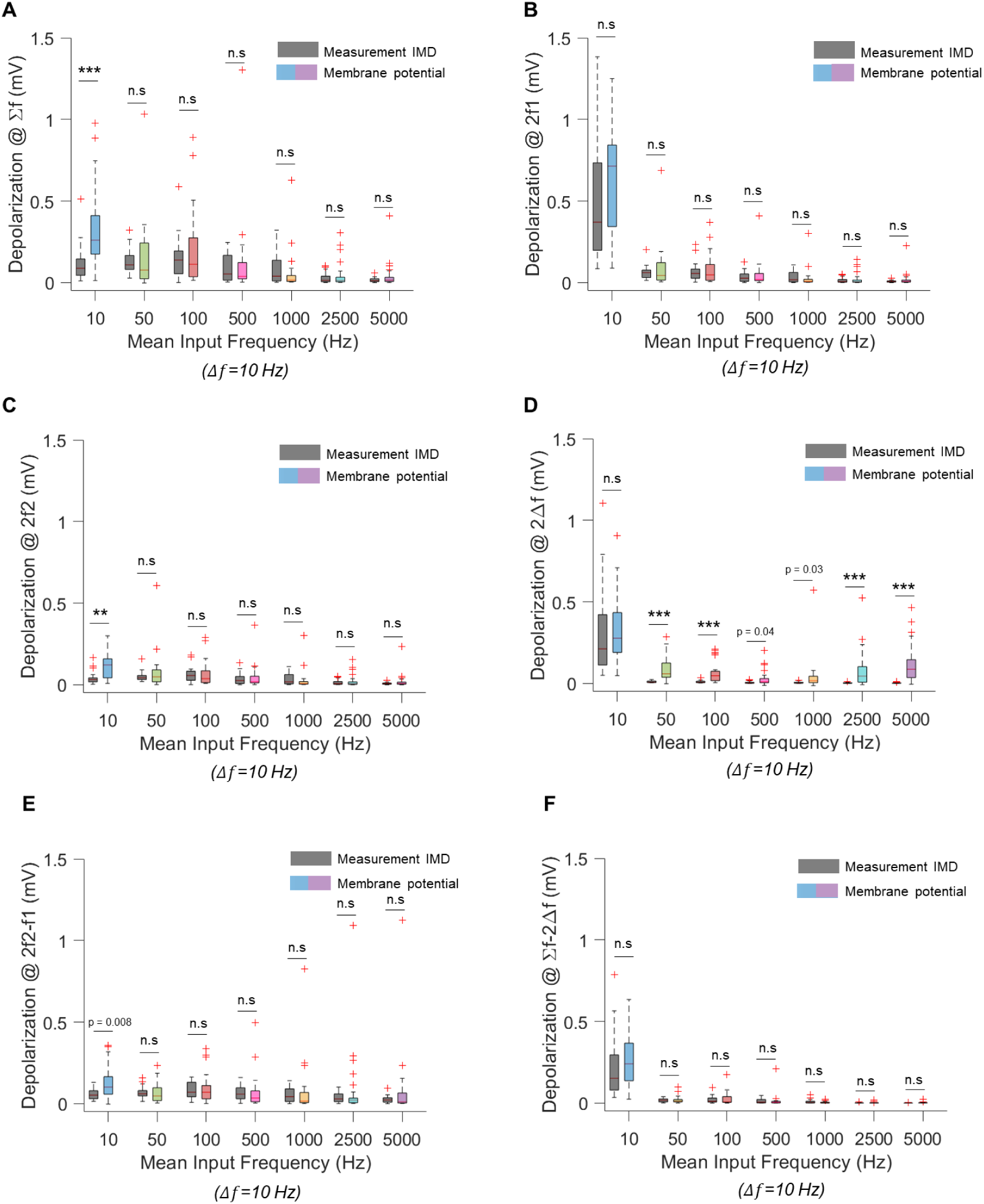
Mixing of exogenous membrane potentials in individual neurons *ex vivo* – additional mixing products. (**A**) Box plot showing RMS amplitude of the induced neural oscillation at *∑f* (‘Membrane potential) vs. the measurements’ intermodulation distortion (IMD) at *∑f* (‘Measurement IMD’) across a range of stimulation frequencies. Individual membrane potential traces were filtered around *∑f* (bandpass filter, cut-off frequencies *∑f* ± 1*Hz*). (**B**) As in (A) but for 2*f*_1_. Individual membrane potential traces were filtered around 2*f*_1_ (bandpass filter, cut-off frequencies 2*f*_1_ ± 1*Hz*). (**C**) As in (A) but for 2*f*_2_. Individual membrane potential traces were filtered around 2*f*_2_ (bandpass filter, cut-off frequencies 2*f*_2_ ± 1*Hz*). (**D**) As in (A) but for 2Δ*f*. Individual membrane potential traces were filtered around 2Δ*f* (bandpass filter, cut-off frequencies 2Δ*f* ± 1*Hz*). (**E**) As in (A) but for 2*f*_2_ – *f*_1_. Individual membrane potential traces were filtered around 2*f*_2_ – *f*_1_ (bandpass filter, cut-off frequencies 2*f*_2_ – *f*_1_ ± _1_*Hz* (**F**) As in (A) but for *∑f* – 2Δ*f*. Individual membrane potential traces were filtered around *∑f* – 2Δ*f* (bandpass filter, cut-off frequencies *∑f* – 2Δ*f* ± 1*Hz*). n (IMD/Membrane potential) = 20/27 (10Hz); 20/21 (50Hz); 25/27 (100Hz); 26/27 (500Hz); 21/25 (1,000Hz); 29/29 (2,500Hz); 28/29 (5,000Hz) recordings/cells. Current densities: 0.41±0.31 (10Hz); 0.30±0.24 (50Hz); 0.34±0.27 (100Hz); 0.34±0.22 (500Hz); 0.33±0.26 (1,000Hz); 0.45±0.34 (2,500Hz); 0.69±0.37 (5,000Hz) mA/mm^2^. For all plots: *, comparisons survived Bonferroni correction (p-value=0.0071). ***, p < 0.0005; **, p<0.005; n.s, non-significant; Wilcoxon rank sum test/two sample t-test. Boxplots: central mark, median; box edges, 25th and 75th percentiles; whiskers, extend up to 1.5x interquartile range box edges; ‘+’, datapoints outside this range.

**Fig. S4.**
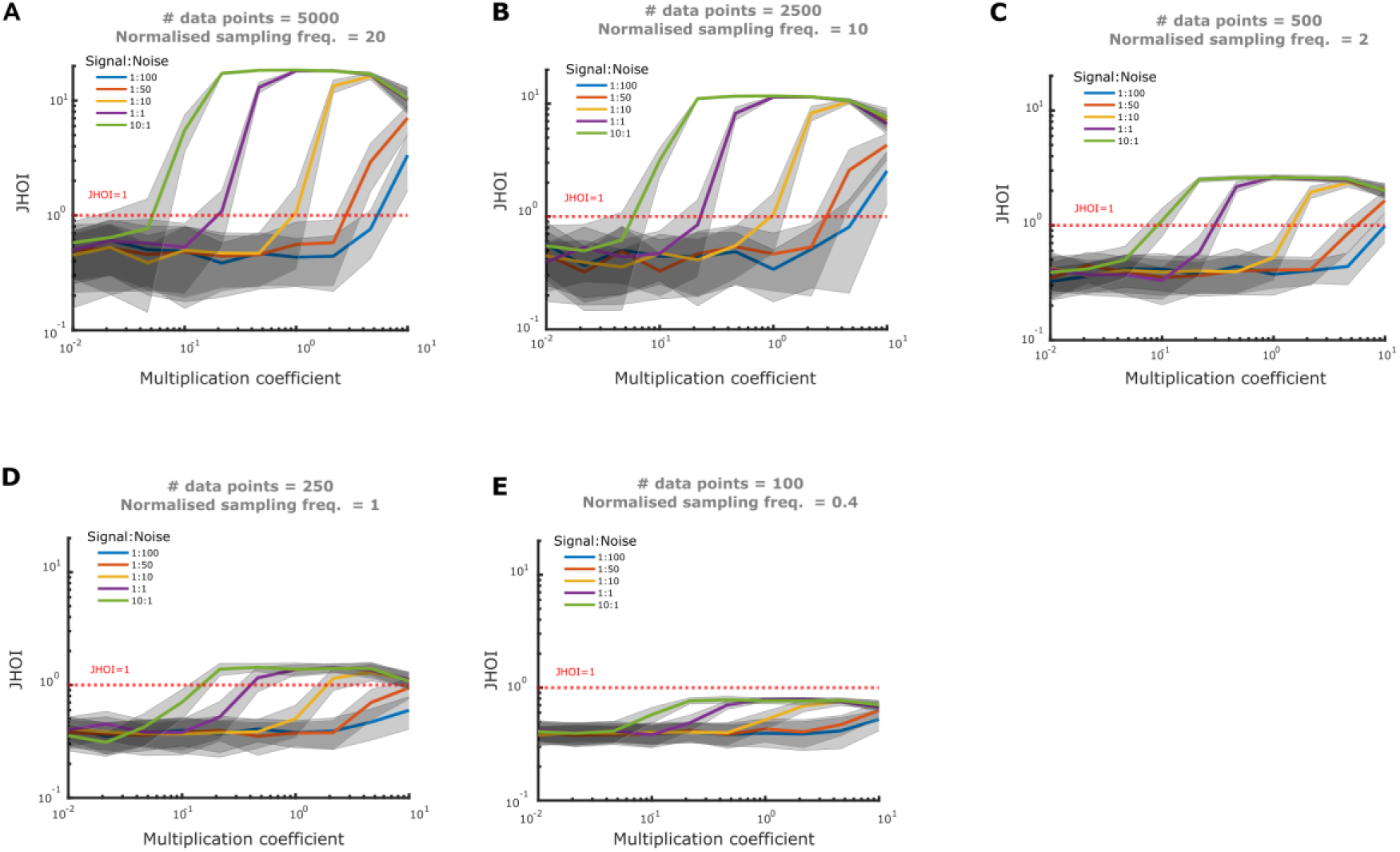
Characterization of JHOI frequency mixing sensitivity with synthetic data. Sensitivity analysis of the JHOI on synthetic data (10 seconds, 500Hz sampling frequency, non-linear mixing (f_1_=7Hz and f_2_=18 Hz). The red dashed line indicates JHOI=1, corresponding to significant frequency mixing relationships (p<0.05). (**A**-**E**) Computing the JHOI for a decreasing number of sampled data points of the instantaneous phase. The normalized sampling frequency is calculated as the sampling frequency (500Hz) divided by the multiplication of the max product frequency (25Hz) and the phase downsampling rate (1, 2, 10, 20, 50). When the normalized sampling frequency is below 1, JHOI is unable to detect significant frequency mixing relationships.

**Fig. S5.**
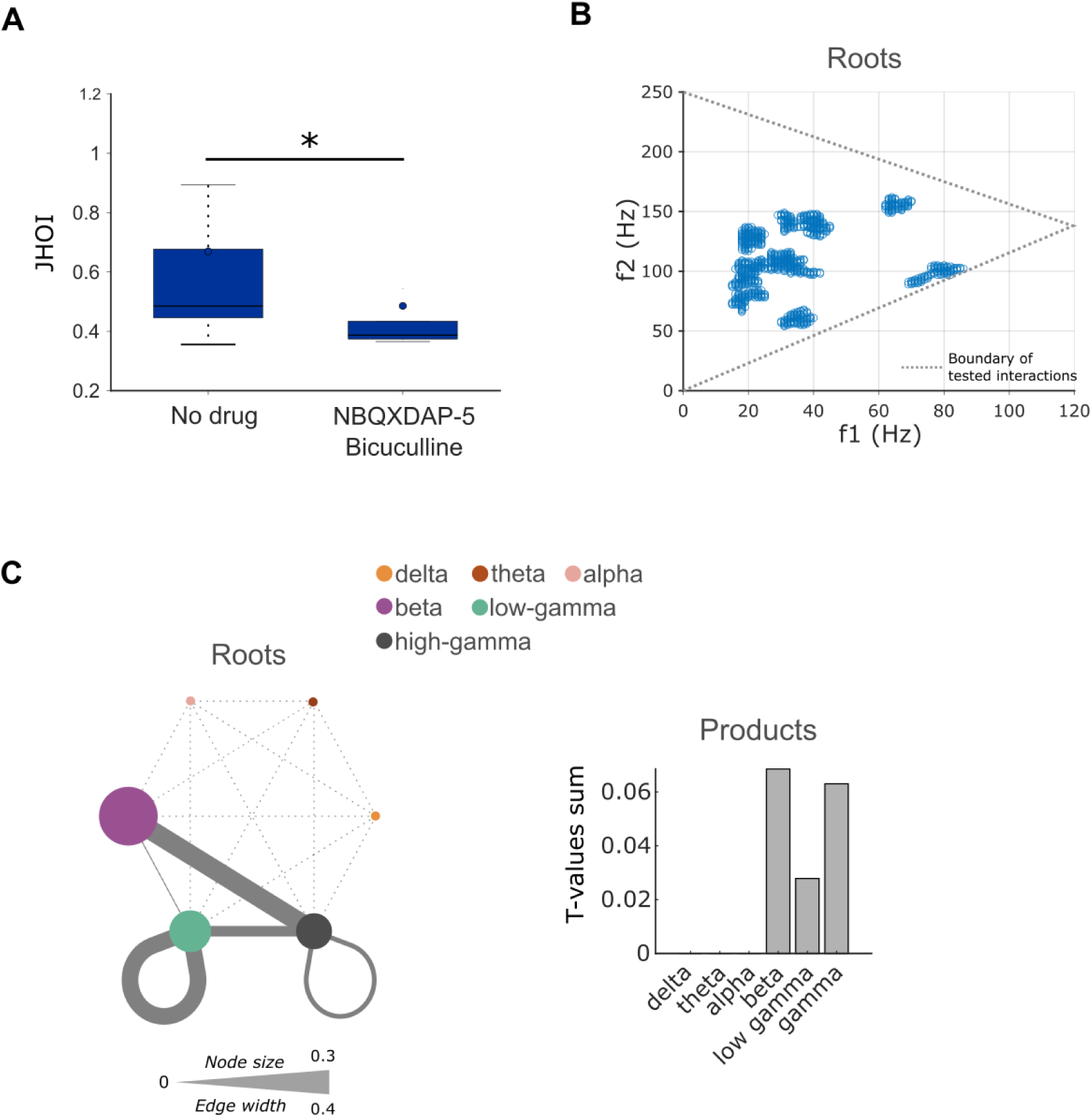
Endogenous membrane potential frequency mixing in individual neurons *ex vivo* – effect of pharmacological blockade. Effect of blockade of NMDA, AMPA, and GABAA ion channel currents ex vivo. n = 8 cells. (**A**) Boxplot of average frequency mixing JHOI across cells. *, p<0.05, paired t-test. (**B**) Frequency-mixing quadruplets reduced at group-level by blockade showing distribution of root frequency clusters. (**C**) same as (B) but showing root frequencies stratified by frequency band and frequency mixing products. Boxplot: central line, median; whiskers, interquartile range; circle, mean; grey dots, outliers.

**Fig. S6.**
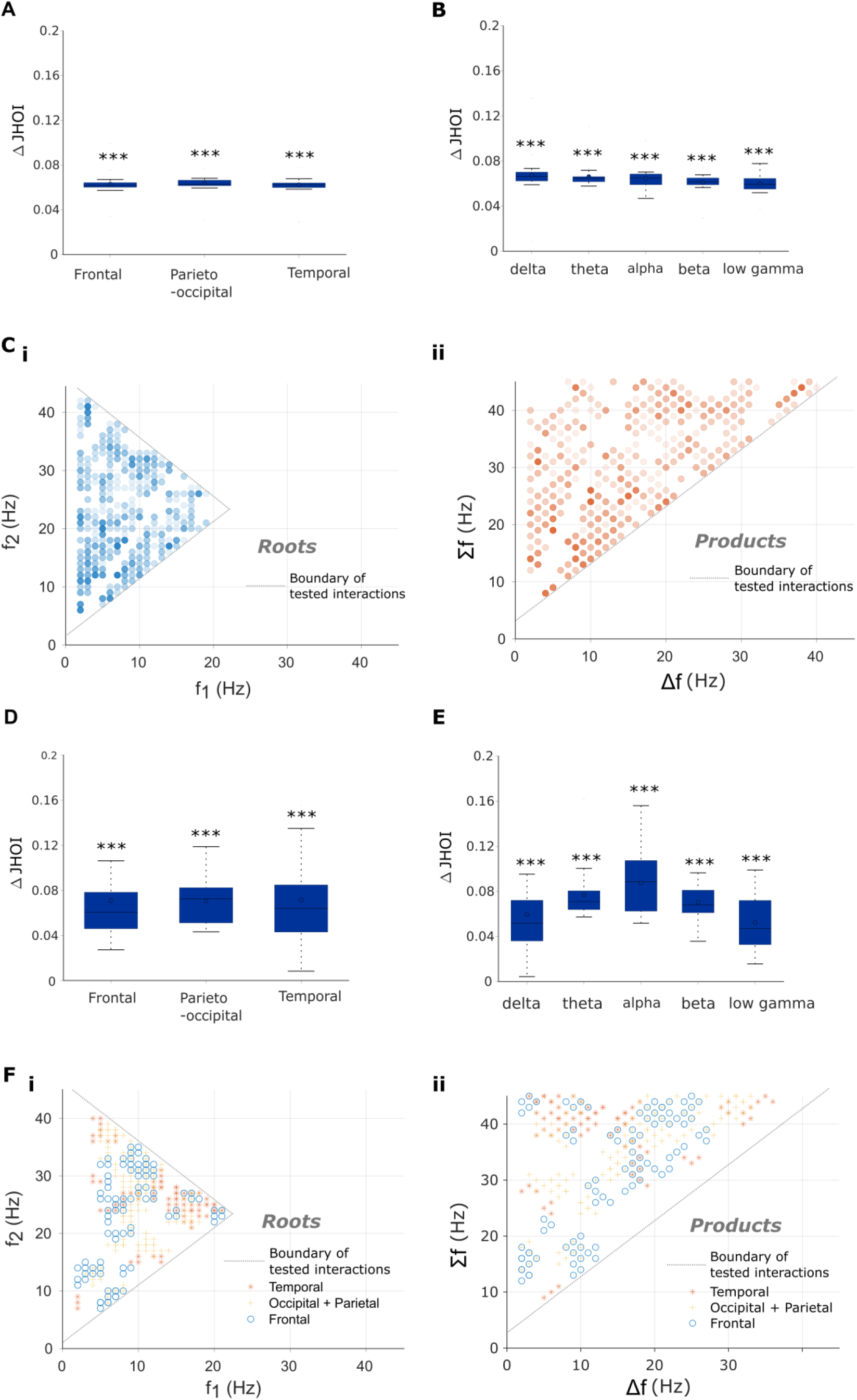
Mixing of endogenous neural network oscillations in the human brain EEG – additional results. (**A**-**C**) Oscillation mixing between brain sites. (**A**) Average frequency mixing JHOI (surrogate subtracted for visualisation) across brain regions. ***, p<0.001, paired t-test against surrogate. (**B**) (i) Average frequency mixing JHOI (surrogate subtracted) across frequency bands. ***, p<0.001, paired t-test against surrogate. (**C**) Frequency quadruplets with significant mixing showing (i) root frequencies and (ii) product frequencies. (**D**-**F**) Local (within brain sites) oscillation mixing. (**D**) As in (**A**) but for local mixing. (**E**) As in (**B**) but for local mixing. (**F**) As (**C**) but for local mixing. Boxplots: central line, median; whiskers, interquartile range.

**Fig. S7.**
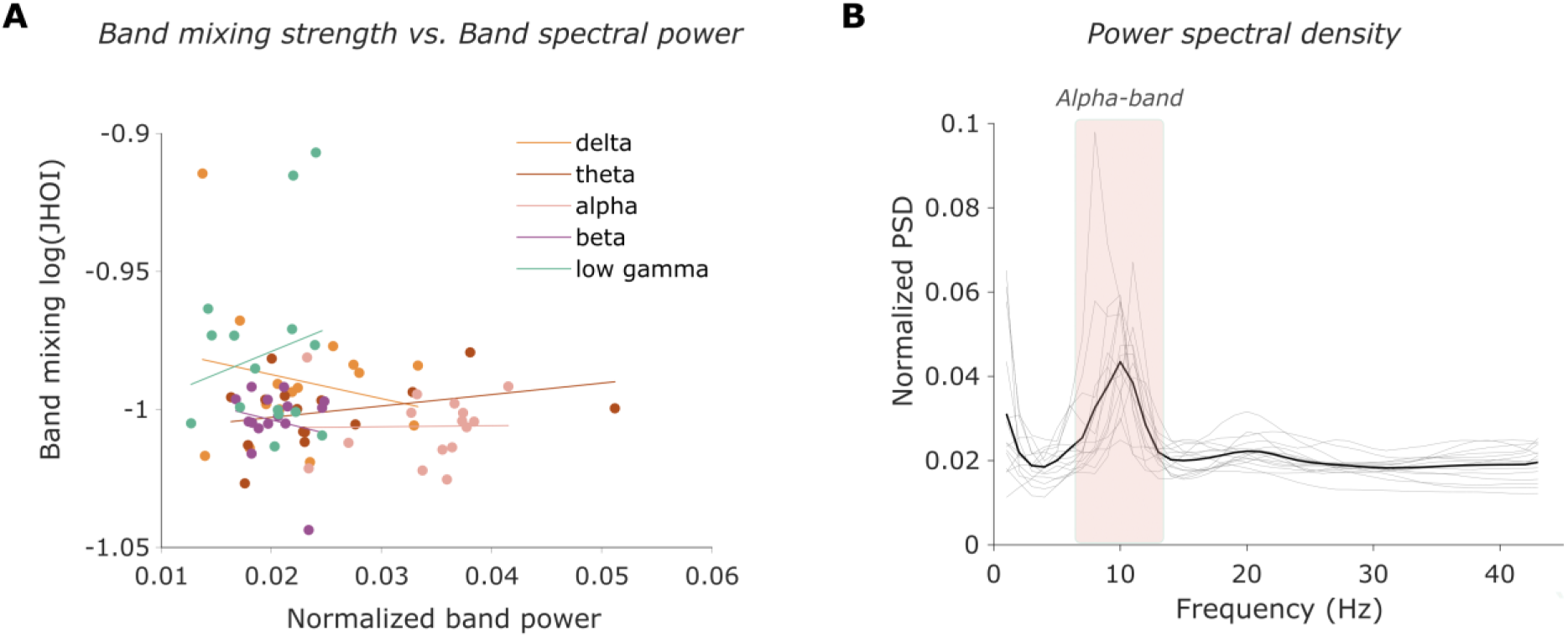
Mixing of endogenous neural network oscillations in the human brain EEG – additional results. (**A**) Participants’ strength of mixing within a band (log JHOI averaged across all inter-site and local mixings) vs power band (normalized to total power). P>0.05 for all correlations. (**B**) EEG power spectral density (PSD) normalized to total power, showing peak at alpha band.

**Table S1.**
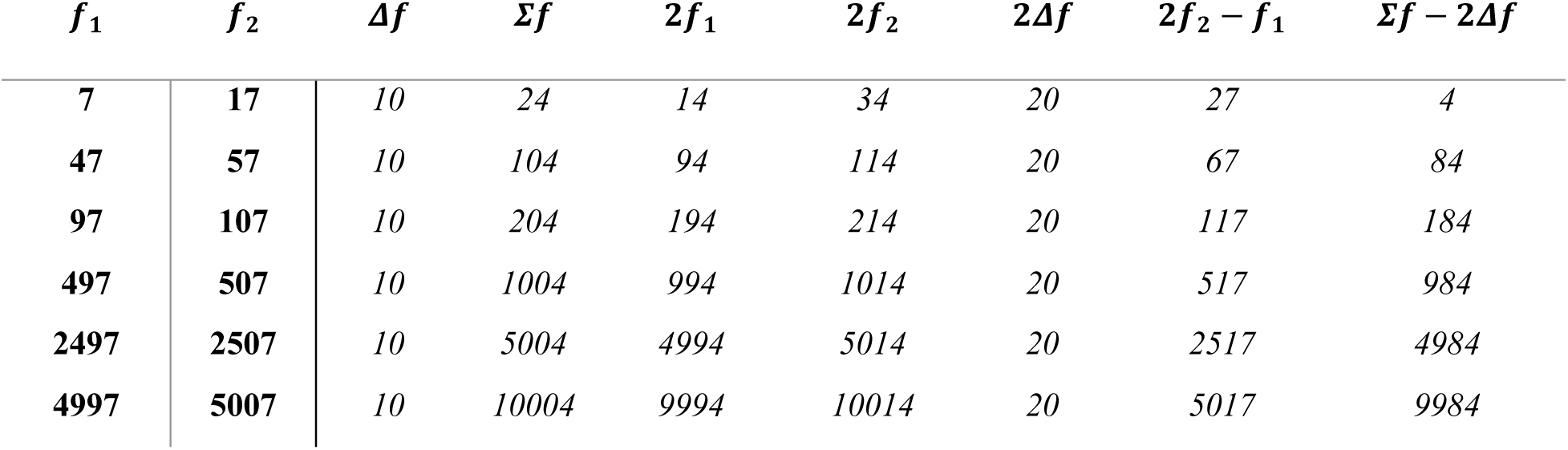
Summary table of applied/input frequencies (bold) and investigated emergent frequencies (italic) for single neuron patch clamp investigation.

**Table S2.** Summary table of statistics for data shown in Figure 1. *x̄* = mean, *σ* = standard deviation. * Indicates significance that survived Bonferroni correction. Bonferroni-corrected p-value = 0.0071, *** p < 0.0005.

| Mean Input Frequency (Hz) | Depolarization (membrane potential) ($\Delta f$) (mV) | | Depolarization (measurement IMD) ($\Delta f$) (mV) | | Two-sample t-test/Wilcoxon rank sum test |
| --- | --- | --- | --- | --- | --- |
| | $\bar{x}$ | $\sigma$ | $\bar{x}$ | $\sigma$ | p |
| 10 | 0.79 | 0.58 | 0.26 | 0.20 | 2.91e-04*** |
| 50 | 0.45 | 0.42 | 0.042 | 0.028 | 1.25e-05*** |
| 100 | 0.34 | 0.30 | 0.038 | 0.026 | 1.62e-07*** |
| 500 | 0.32 | 0.37 | 0.017 | 0.011 | 6.67e-08*** |
| 1000 | 0.36 | 0.55 | 0.015 | 0.011 | 1.53e-07*** |
| 2500 | 0.55 | 0.50 | 0.011 | 0.011 | 8.72e-09*** |
| 5000 | 0.76 | 0.51 | 0.011 | 0.0091 | 1.069e-10*** |

**Table S3.** Summary table of statistics for data shown in Figure 1 cont. *x̄* = mean, *σ* = standard deviation. * Indicates significance that survived Bonferroni correction. Bonferroni-corrected p-value = 0.0021, **, p < 0.005; ***, p < 0.0005.

| Mean Input Frequency (Hz) | Normalized Threshold (a.u.) |  | Post-hoc paired t-test |  |  |  |  |  |  |
| --- | --- | --- | --- | --- | --- | --- | --- | --- | --- |
| | $\bar{x}$ | $\sigma$ | p (vs. 10Hz) | p (vs. 50Hz) | p (vs. 100Hz) | p (vs. 500Hz) | p (vs. 1000Hz) | p (vs. 2500Hz) | p (vs. 5000Hz) |
| 10 | 1.19 | 0.39 | - | 0.23 | 0.30 | 0.58 | 0.48 | 0.018 | 4.43e-05*** |
| 50 | 1.06 | 0.40 | 0.23 | - | 0.77 | 0.86 | 0.21 | 0.0022** | 2.94e-06*** |
| 100 | 1.09 | 0.47 | 0.30 | 0.77 | - | 0.98 | 0.24 | 0.0045 | 5.44e-06*** |
| 500 | 1.09 | 0.84 | 0.58 | 0.86 | 0.98 | - | 0.096 | 5.56e-05*** | 8.64e-07*** |
| 1000 | 1.30 | 0.77 | 0.48 | 0.21 | 0.24 | 0.0096 | - | 0.018 | 3.58e-07*** |
| 2500 | 1.72 | 0.87 | 0.018 | 0.0022** | 0.0045 | 5.56e-05*** | 0.018 | - | 4.58e-04*** |
| 5000 | 2.82 | 1.31 | 4.43e-05*** | 2.94e-06*** | 5.44e-06*** | 8.64e-07*** | 3.58e-07*** | 4.58e-04*** | - |

**Table S4.**
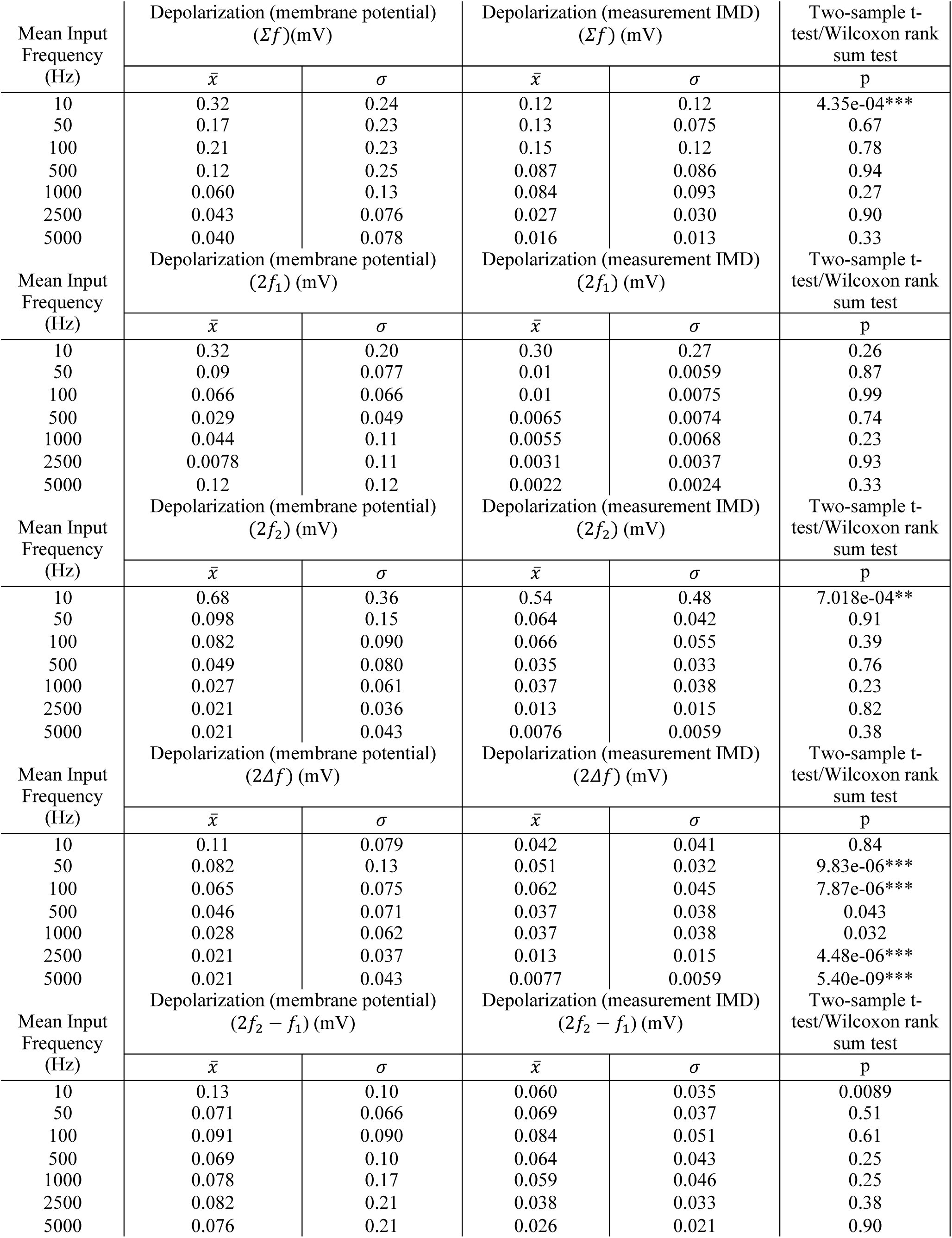
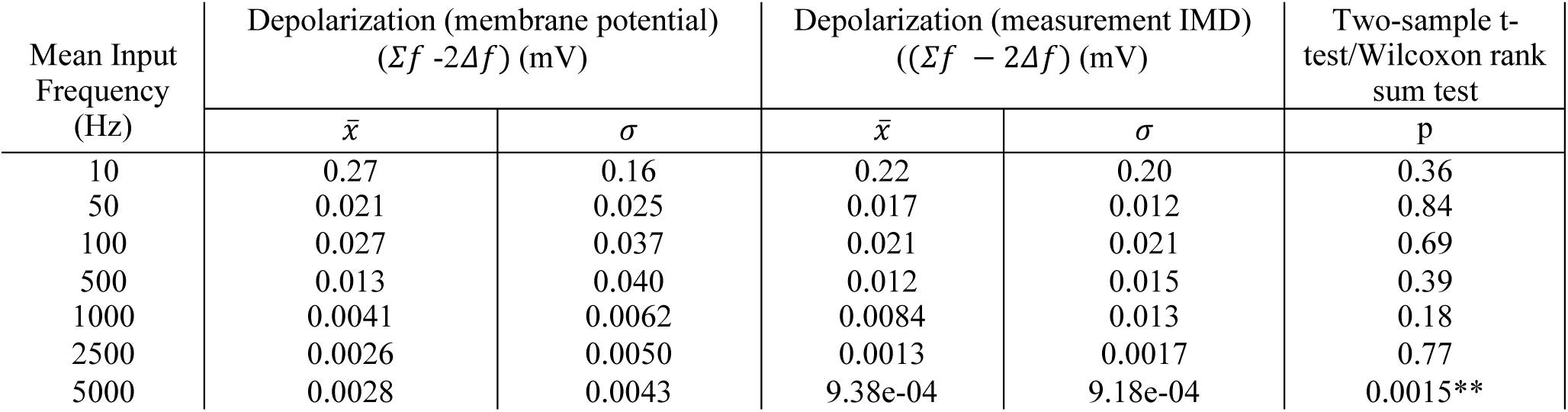
Summary table of statistics for data shown in Figure S3. *x̄* = mean, *σ* = standard deviation. * Indicates significance that survived Bonferroni correction. Bonferroni-corrected p-value = 0.0071, **, p < 0.005.

**Table S5.**
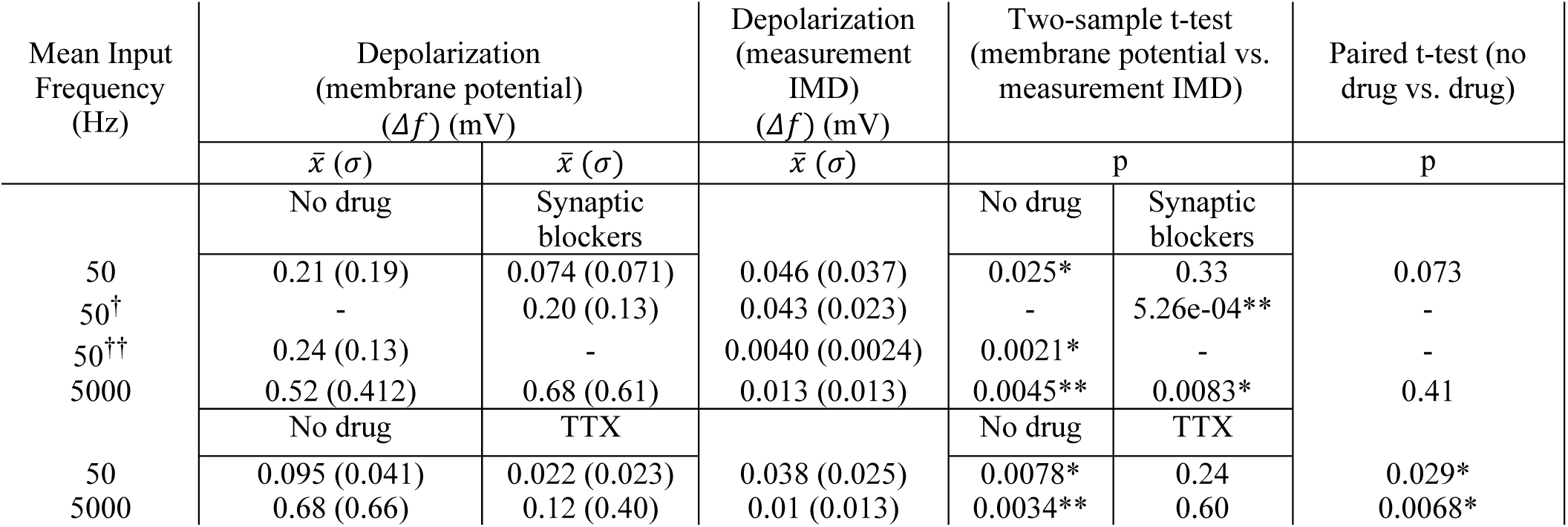
Summary table of statistics for data shown in Figure 2. *x̄* = mean, *σ* = standard deviation. * Indicates significance that survived Bonferroni correction if required. Bonferroni-corrected p-value for two sample t-test = 0.025, *, p < 0.05; **, p < 0.005. † indicates higher current density experiments. †† indicates intracellular stimulation experiment.

